# Simulation of Pedigree vs. fully-informative marker based relationships matrices in a Loblolly Pine breeding population

**DOI:** 10.1101/2021.11.30.468863

**Authors:** Adam R Festa, Ross Whetten

## Abstract

Computer simulations of breeding strategies are an essential resource for tree breeders because they allow exploratory analyses into potential long-term impacts on genetic gain and inbreeding consequences without bearing the cost, time, or resource requirements of field experiments. Previous work has modeled the potential long-term implications on inbreeding and genetic gain using random mating and phenotypic selection. Reduction in sequencing costs has enabled the use of DNA marker-based relationship matrices in addition to or in place of pedigree-based allele sharing estimates; this has been shown to provide a significant increase in the accuracy of progeny breeding value prediction. A potential pitfall of genomic selection using genetic relationship matrices is increased coancestry among selections, leading to the accumulation of deleterious alleles and inbreeding depression. We used simulation to compare the relative genetic gain and risk of inbreeding depression within a breeding program similar to loblolly pine, utilizing pedigree-based or marker-based relationships over ten generations. We saw a faster rate of purging deleterious alleles when using a genomic relationship matrix based on markers that track identity-by-descent of segments of the genome. Additionally, we observed an increase in the rate of genetic gain when using a genomic relationship matrix instead of a pedigree-based relationship matrix. While the genetic variance of populations decreased more rapidly when using genomic-based relationship matrices as opposed to pedigree-based, there appeared to be no long-term consequences on the accumulation of deleterious alleles within the simulated breeding strategy.

## Introduction

Loblolly pine (*Pinus taeda*) is one of the most important commercial tree species in the United States, with over 700 million seedlings planted annually [1]. Over the past 60 years, Tree Improvement Cooperatives (TICs) have formed to help meet the growing need for increased forest productivity and resource management. The main objective of these Cooperatives is to develop future generations of genetically improved loblolly pine that will provide more wood on less land in less time. Given a large amount of genetic variation in loblolly pine, the ability to make a significant amount of genetic gain on traits such as height, diameter, and disease resistance will help meet this objective [2]. However, TICs face a trade-off in the rate at which genetic gain can be obtained by balancing selection intensity with genetic similarity among future prospective selections. This trade-off occurs because breeding individuals with increased genetic similarity can lead to inbreeding depression and cause a severe reduction in the phenotypes of high economic importance [3].

Inbreeding depression can be defined as a decline in mean phenotype due to mating among individuals with similar coancestry, resulting in increased homozygosity within populations [4]. The decline in phenotype can be caused by the effects of loci that harbor recessive deleterious alleles. While multiple theories exist for the genetic basis of traits displaying inbreeding depression, they all incorporate the concept that alleles involved are under some effect of dominance. In contrast, the average effect associated with alleles in the progeny is independent of the parents’ inbreeding level for a purely additive trait. Because of this, impacts of inbreeding depression within a breeding program may potentially be minimized by identifying and purging deleterious alleles or utilizing mating designs that are aimed at controlling the increase of homozygosity within breeding populations [5].

The effects of inbreeding depression within conifers have been extensively studied and shown to have consequences in seed production and multiple phenotypic traits [5, 9]. The prevalence of inbreeding depression in loblolly pine has motivated studies to explore the mean number of these alleles per individual within a given population [6]. Two independent studies have been conducted within loblolly pine aimed at quantifying the number of recessive lethal alleles. Franklin (1972) analyzed progeny from the self-pollination of 116 parent trees and reported that lethal equivalents followed a relatively normal distribution with a mean of 8.5 [7]. The second study by Remington et al. (2000) conducted whole-genome characterization of embryonic inbreeding depression in progeny from self-pollination of an elite loblolly pine tree and found roughly 3-6 lethal alleles [8]. Recently, Ford et al. analyzed twenty loblolly pine parent trees mated to have progeny with varying degrees of inbreeding coefficients. At the highest inbreeding coefficient within the population analyzed, they reported a 21% and 33% reduction in height and stem volume, respectively [10]. Based on the literature on the effects and possible long-term impacts of inbreeding, it is essential that TICs properly manage the accumulation of deleterious alleles within their breeding programs.

Breeding strategies for loblolly pine have evolved [11,12]. The North Carolina State University cooperative tree improvement program currently uses a rolling front strategy. New parents are crossed every year, new progeny tests are established, and pre-existing progeny tests are measured to update family breeding values. This strategy has been shown to increase genetic gain up to 25-30% relative to a discrete generation strategy, allowing for more phenotypic gain to be made each year [13]. For selections and mating designs, inbreeding management among individuals within the North Carolina State University TIC is currently handled using the computer program Mate Select, which employs the evolutionary differential algorithm in utilizing information obtained from progeny tests to develop a mating design, taking both performance and amount of relatedness into account [14]. The accuracy of relatedness estimations between selections is an important consideration when producing future generations of loblolly pine. Not only can the consequences of inbreeding depression be severe, but given the long generation interval, it could take a substantial amount of time to overcome.

The method currently used to estimate relatedness among individuals to make selections is based on average levels of allele sharing extracted from pedigree data. However, unlike most other breeding programs, tree breeding programs are in their infancy, typically no more than 3 or 4 generations deep with relatively weak pedigrees and connectedness among progeny test datasets [15]. The idea of using genomic markers for estimating relatedness has existed for multiple decades [16]. More recently, the reduction in the cost of obtaining genomic markers has inspired novel methods of estimating genomic relationship matrices and implementing genomic selection [17, 18]. Genomic selection is the process of selecting new breeding parents based upon integrating phenotypic and shared genotypic information instead of relying on phenotype and assumed pedigree allele sharing [19]. Provided with access to these genomic resources, the use of DNA marker-based relationship matrices, in addition to or in place of pedigree-based allele sharing estimates, has shown to provide a significant increase in the accuracy of progeny breeding value prediction [20-25]

Given the success reported in breeding programs for other species, it is intriguing to consider how utilizing marker-based versus pedigree-based relationship matrices in loblolly pine breeding selection strategies would impact long-term inbreeding depression. Additionally, it would be beneficial in exploring future selection regimes to have the ability to assess breeding strategies for multiple generations and evaluate potential long-term genetic gain or risk of inbreeding. A computer simulation allows the flexibility to address these questions without bearing the cost of field experiments and limits on population size or generation length.

Computer simulations have been developed to explore the consequences of different breeding strategies in tree programs over the past 20 years and were shown to be a useful tool [26-34]. Few simulation studies have been published for loblolly pine, and most of those are generally aimed at topics other than genomic selection [32-34]. While many breeding simulation programs exist, currently available programs cannot assess the genetic value of progeny from multiple generations with a user-defined genetic architecture for traits to be simulated. Our purpose for creating a simulation program is to enable an exploratory analysis of the potential genetic architecture for phenotypic traits, assessment of potential long-term impacts of different selection strategies, and the ability to predict possible long-term inbreeding consequences. This simulation program will be a helpful resource when comparing multiple long-term breeding and selection strategies for their ability to sustain long-term genetic and economic gain.

The overall objective of this study is to compare the relative genetic gain within a breeding program similar to that of loblolly pine, utilizing pedigree-based or marker-based relationships. Developing a computer simulation program that can simulate multiple generations utilizing marker and phenotypic data to predict progeny genetic values and phenotypes for a given genetic architecture will aid in accomplishing this objective. The first goal of this study is to create a breeding simulator and validate the reliability by creating a genetic linkage map using simulated genotype data for an F2 population; the second goal is to explore the possible parameter space to match loblolly pine’s current proposed genetic architecture to replicate results from past field experiments; and the final goal is to test the impact of using either fully informative marker-based or pedigree-based relationship matrices on inbreeding depression and the expected amount of genetic gain utilizing an assortative mating design. We also explore the relationship between the number of parents and progeny per cross with the impact on genetic gain across a multi-generation breeding strategy.

## Methods

### Creation of a forward population simulator

The simulator package was written in R and relies on various other R packages used to construct pedigree relationship matrices and evaluate inbreeding statistics, such as the pedigree package [35-40]. All source code, functions, and descriptions are in the appendix. In general, we generated multiple functions that helped to facilitate the simulation of a forward breeding population based on the user’s input, including:

- Creation of a genetic map (create_genetic_map)
- Simulation of a founder population (create_founder_parents)
- Creation of a cross design schema (create_cross_design)
- Simulation of parental crosses (make_crosses)
- Estimation of genotypic and phenotypic values (calc_tgv and sim_phenos)
- Extraction of selections for next-generation (extract_selections)
- Simulation of Open Pollinated testing (OP_testing)

We utilized each of these for all forward population simulations and changed input parameters as described below. Additionally, we describe the simulator methods as needed, and further explanation may be found in the appendix.

### Simulation of a Linkage Map

A genetic linkage map was produced to validate that the simulator could create the correct number of linkage groups and order of markers. The most efficient linkage map algorithm, mstmap, does not work with outbred populations such as loblolly pine; therefore, a 2^nd^ generation population was generated and used as the input. We set the create_genetic_map function to simulate a genetic architecture with 12 linkage groups (LG), 120 markers, and a map length equal to roughly1800cM. Each LG was assigned ten markers, equally spaced. We simulated two unrelated inbred parents with different homozygous allelic states at each marker locus. A single F1 individual was selfed to generate 300 F2 progeny. Within the ASMap R package, this type of population is referred to as RIL2 [41]. The marker matrix returned from the calc_tgv function was converted from 0,1,2 output to A, B, X and used as the input to the R package ASMap to generate a linkage map. The resulting linkage map was compared to that expected based on input parameters. [41].

### Replication of past field experiments

We adjusted inputs of the simulator to identify allelic and dominance values that represent loblolly pine inbreeding characteristics similar to those previously found in the literature. The create_genetic_map parameters were set to generate 12 LGs with a genetic map size equal to roughly 1800 cM [42]. A total of 2760 loci were distributed evenly across the 12 LGs so that each had 230 loci. The 2760 loci were divided into three separate categories: *random* QTL, *SNP* QTL, and markers. *Random* QTL (QTL_r_) represents random genetic noise caused by events such as epistasis or regulatory interactions. *SNP* QTL (QTLSNP) represents loci that can show a dominance effect, and markers are loci that may be used for genomic selection and estimation of progeny relatedness. The number of loci assigned to each of the three categories was 680 QTL_r_, 1960 QTLSNP, & 120 markers. The simulated markers were not used in this analysis, but were included so the simulator parameters would be as similar as possible to those used later in the breeding population simulations. The other two types of loci, QTL_r_ and QTLSNP, were randomly assigned to positions on the linkage groups. The recombination rate of a particular locus was determined by utilizing the genetic distance between each pair of adjacent loci with the Haldane function [43]. Minor allele frequencies (MAF’s) for QTLSNP were sampled from a uniform distribution ranging from 0.01 to 0.02, because this category was used to simulate deleterious alleles that contribute to inbreeding depression, and such alleles are believed to be rare in the pine population. QTL_r_ MAF’s were sampled from a beta distribution with alpha and beta set to equal 0.4. Two traits (straightness and volume) were evaluated at four inbreeding coefficient levels (0, 0.125, 0.25, 0.5). The dominance and major/minor allele values assigned for each of the traits are reported in Table 1.

**Table 1.** Simulator input values of dominance coefficient, major and minor alleles used to model inbreeding depression in two traits (volume and straightness).

| Trait | Dominance Coefficient | Major Allele Value | Minor Allele |
| --- | --- | --- | --- |
| Volume | 1 | 1 | -100 |
| Straightness | .5 | 1 | -1 |

An initial founder set of 50 unrelated parents was generated utilizing the genetic map to create a population of individuals which contain unrelated, half-sib, and full-sib individuals. QTL_r_ values, which are assigned to the 680 random QTL on the genetic map, were sampled for each parent from a normal distribution ranging from -1 to +1 with a mean equal to 0. Genetic effects at random QTL are completely additive, but the genetic effects at QTLSNP loci had a dominance component. The total genetic value due to QTLSNP loci within an individual was calculated as:

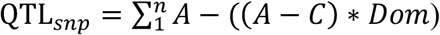

Where (*A*) represents the value of the major allele, (*C*) represents the value of the minor allele, and *Dom* is the dominance coefficient applied to that locus. The inbreeding experiment was simulated starting with unrelated parents, so each parent was assigned heterozygous recessive deleterious alleles at different loci. Assigning alleles in this way means that, in the absence of inbreeding, there will be no chance that crossing any two parents will result in the presence of deleterious alleles in the homozygous state. To mimic the number of deleterious alleles cited in the literature, the QTLSNP were assigned to parents to provide an average of seven recessive alleles per parent. Once founder parent alleles are assigned, the genetic value of a given individual is calculated as:

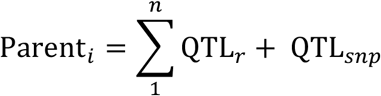

A user-specified cross-design file was supplied to make 40 crosses among the parents with two progeny each. A single selection was made within each family based upon the phenotype. The resulting 40 selections were then mated so that ten parents were crossed to two unrelated individuals, two half-sibs, two full sibs, and each of the ten was selfed. This mating design is based on that used by Ford et al (2015) as shown in Figure 1 below. Fifty progeny were created for each cross.

**Figure 1.**
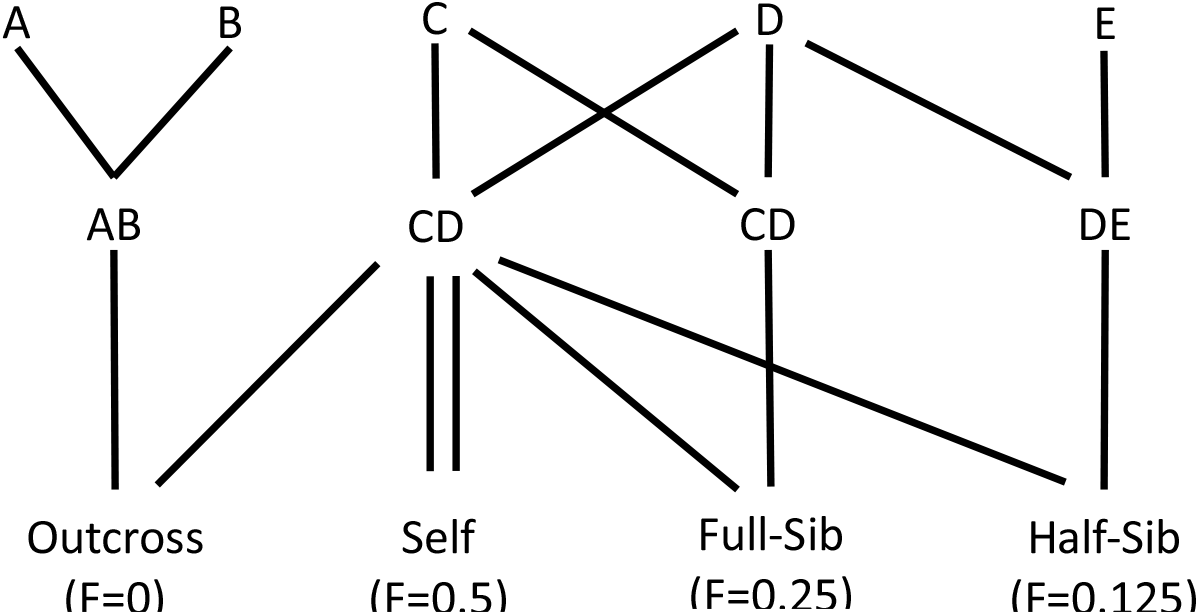
Simulated cross design demonstrating the production of the four levels of inbreeding. Parents A, B, C, D, and E are initial founder parents. Parent CD1 was crossed with two unrelated individuals (e.g., AB), two full-sibs (e.g., CD2), two half-sibs (e.g., DE), and selfed (CD1 × CD1) to produce each family.

Simulation of phenotypes was done by adding random noise sampled from a normal distribution with a mean and standard deviation of:

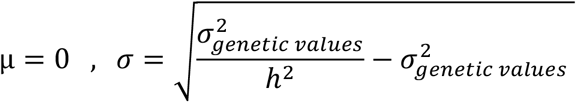

Phenotypes were assigned to the founder population so that individual tree heritability was roughly h^2^=0.3. All subsequent progeny phenotypes were estimated using a standard deviation equal to that within the founder population. Given that phenotypes are determined by adding random environmental noise to genetic values, progeny phenotypes returned from the simulator were transformed into values representing those typically collected in the field. Volume phenotypes were converted to a range from 20-140 by applying the following formula

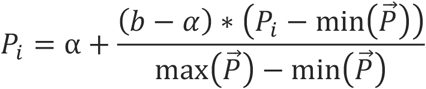

where *P*_*i*_ is the phenotype of an individual, α is the minimum range, *b* is the maximum range, and 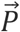 is the vector of all progeny phenotypes. The following linear mixed model was used to analyze volume. Inbreeding level was treated as a fixed effect, and parents were considered random effects.

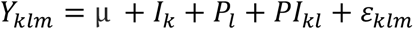

where *Y*_*klm*_ is the response variable of the *m*th tree of the *l*th parent at the *k*th inbreeding level, μ is the overall mean, *I*_*k*_ is the fixed *k*th inbreeding level effect (*k* = 4), *P*_*l*_ is the random *l*th parent effect (*l* = 10) with expectations NID ∼(0, σ_1_^2^*i*), *PI*_*lk*_ is the random interaction of the *l*th parent with the *k*th inbreeding level with expectations NID ∼(0, σ*lk*^2^), and ε_*klm*_ is the random error term with expectations NID ∼(0, σ*e*^2^)

The ratios of variance components to their standard errors (Est/SE) were computed for the random terms. With sufficient factor levels, the ratio can be interpreted as an asymptotic approximation of the *z* distribution, which can be used for a hypothesis test for the effect of the null random terms [44,45].

The lmer function within lme4 software and Tukey’s means separation was used to obtain the significant differences among the four levels of inbreeding [46]. Percent inbreeding depression was calculated for each trait and defined as the percent reduction in the mean of the trait relative to the mean of progeny from unrelated crosses. A linear model was used to estimate the effect that starting number of deleterious alleles had on the phenotype of the selfed parent.

Stem straightness was converted to a binary trait by specifying 1 for individuals with the above mean phenotype and 0 for those below the mean phenotype. The probability of straightness was modeled by fitting a generalized linear mixed model in the glmr procedure [46]. The generalized linear mixed model and expectations are as follows

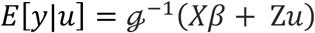

where **y** is (*nx1*) response vector, **X** is (*n* × *p*) design matrix of rank *k* for the (*p* × *1*) fixed effects **β, Z** is (*n* × *q*) design matrix for the (*q* × *1*) random effects, **u**. Random effects are assumed to be normally distributed with a mean equal to 0 and variance matrix **G**. The logit link function, *g* = ln(μ_*i*_(1 − μ_*i*_)) was used to model the binomially distributed response variable, where *ln* is the natural logarithm, and μ_*i*_ is the mean incidence. Therefore, the inverse link function was used to obtain the probability of above-average straightness as *g*^−1^ = *e*^*n*^ = 1/(1 + *e*^*n*^).

### Forward population breeding strategies

The goal of simulating forward breeding strategies is to compare the rate of genetic gain and the risk of inbreeding depression. To mimic the inbreeding depression in loblolly pine, we utilized parameters identified from the simulation of an inbreeding experiment that produced results similar to those observed in field experiments. The strategies compared included making selections within-family utilizing relationship matrices based either on pedigree information or fully informative genetic markers, as well as breeding populations of different sizes.

Founder parents were simulated to have multiple possible haplotype combinations or “fully-informative” markers, meaning that every haplotype within each parent is unique, rather than simulating bi-allelic markers. Simulating markers in this way allows identity by descent to be unambiguously tracked across each generation. A unique set of markers was assigned to each parent using two alphabetical letters for each allele to simulate fully informative markers for calculation of marker-based relationship matrices. Capital letters represent the major allele, and lower-case letters represent the minor allele. All marker loci in the founder populations were heterozygous at each locus so that haplotype allele sharing could be tracked across generations (Figure 2).

**Figure 2.**
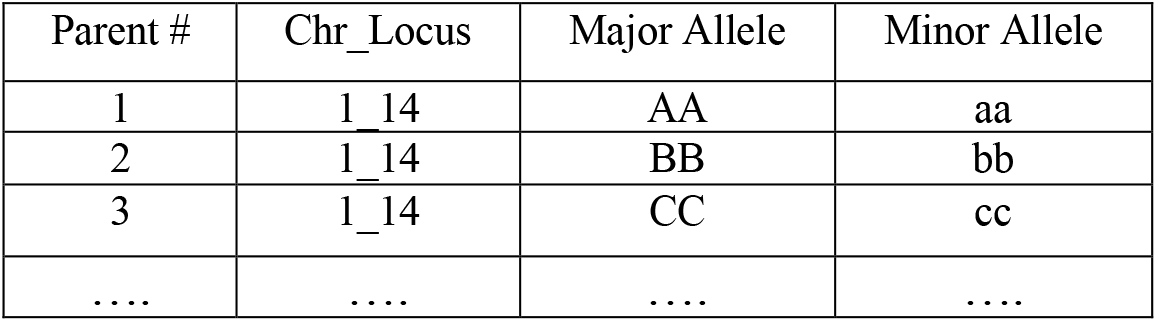
Example of generating unique multi-allelic haplotype markers for each parent at a given marker locus

Ten out of the total 230 loci were sampled for each LG and designated as markers. Selection of marker positions was made by identifying loci closest to 10 equally spaced sites within each LG. The characteristics and assignment of markers utilized within this simulation are made with the assumption that any breeding program in the genomic era has the opportunity to select equally spaced fully-informative markers rather than just using a random sample of bi-allelic marker loci.

A total of 300 genetic maps were simulated to generate 300 unique base populations containing 280 parents. Once a founder population was established, it was separated into three groups comprising the top 64, 140, or all 280 parents based on their respective breeding values estimated from an open-pollinated mating design. An open-pollinated mating design was accomplished by crossing each of 280 parents to the other 279 parents, with a single progeny produced from each cross, and simulating phenotypes for the progeny based on the combination of genetic value and random noise as described previously. The mean of simulated phenotypes from progeny was used to rank parents from best to worst – there was no need to use linear modeling of these phenotypes because they were not simulated to include test, rep, or block effects that modeling could account for.

Each parent base population size was crossed using an assortative mating strategy with a coancestry threshold. An overview of this breeding strategy is in the Figure 3 below. Founder parents were first sorted according to breeding value and then separated into four bins. A starting coancestry threshold of 0.01 was applied to the pedigree relationship matrix, and any two pairs which had a relatedness less than that could potentially mate. The coancestry threshold was increased by 0.005 until the top two quartiles of individuals could be mated to one another. The 64 and 140 parent base populations were set to have 60 progeny for each cross. To investigate the impact of increasing the base population size while decreasing progeny, the parent base population of 280 parents had 30 progeny per cross. Individuals in the top quartile were crossed four times, those in the second quartile were crossed three times, and those in the bottom two quartiles were crossed twice each. This mating design aims to both provide opportunities to identify elite progeny from crosses among the best parents and improve the average performance of the worst parents by crossing them to better-performing individuals.

**Figure 3.**
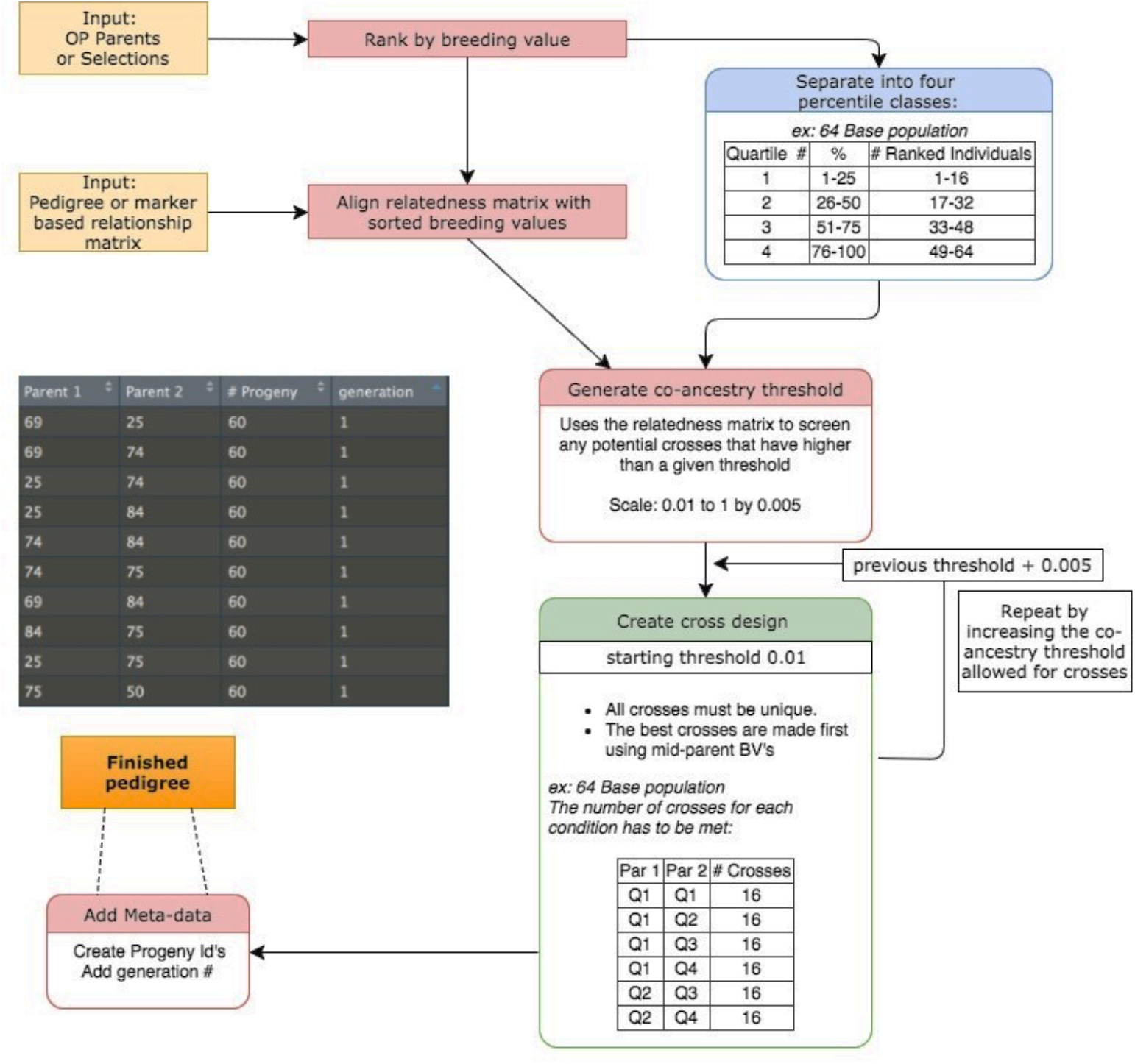
Schematic of generating a cross design matrix for each generation in the 64 base population by applying an assortative mating design with coancestry threshold

For each of the base populations, after making crosses, genetic values and phenotypes were estimated, and selections were made using the best linear unbiased prediction (BLUP) method with either a pedigree-based relationship matrix (ABLUP) or genomic relationship matrix (GBLUP). Among-family selections were made to reduce the initial 96, 210, 420 crosses to the top 64,140, 280 individuals respectively, by ranking family mean BLUP values estimated from progeny. Within-family selections were made by selecting the highest individual progeny BLUP. This resulted in either 64, 140, or 280 selections from each generation used as parents for the next generation. Genetic gain, the average number of deleterious alleles, and genotypic variance of selections were analyzed across ten generations with the following formula

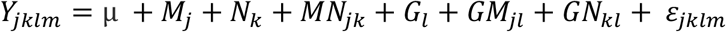

where *Y*_*jklm*_ is the response variable of the *m*th population at the *l*th generation with the *k*th base population number applying the *j*th method, μ is the overall mean, *M*_*j*_ is the fixed jth method (j=2), *N*_*k*_ is the fixed number of individuals within the base population (k=3), *MN*_*jk*_ is the fixed interaction of the jth method with kth base population size, *G*_*l*_ is the random lth generation effect (l=10) with expectations NID ∼(0, σ_1_^2^*i*), *GM*_*jl*_ is the random lth generation interaction with the jth method applied NID ∼(0, σ_jl_^2^), *GN*_*kl*_ is the random lth generation interaction with the number of individuals in the base population NID ∼(0, σ_kl_^2^), and ε _*jklm*_ is the random error term with expectations NID ∼(0, σ*e*^2^).

The ratios of variance components to their standard errors (Est/SE) were computed for the random terms. Tukey’s means separation was used to obtain the significant differences among the two methods for genetic variance across each generation.

## Results

### Linkage Map

The genetic linkage map produced from ASMap contained 12 segregating LGs and a total map length equal to 1886.84 cM. (Figure 4) The size of linkage groups ranged from 144.25 – 170.52cM, and each had ten markers. The mean distance between markers was equal to 17.47cM and a standard deviation of 2.28cM. For comparison, the genetic map length returned from the create_genetic_map function was 1875.49cM and linkage group sizes ranged from 103.9cM - 183.8cM. The mean distance between markers was 15.46 cM with a standard deviation of 1.99cM. All markers in the linkage map were in the same order as the simulated genetic map.

**Figure 4.**
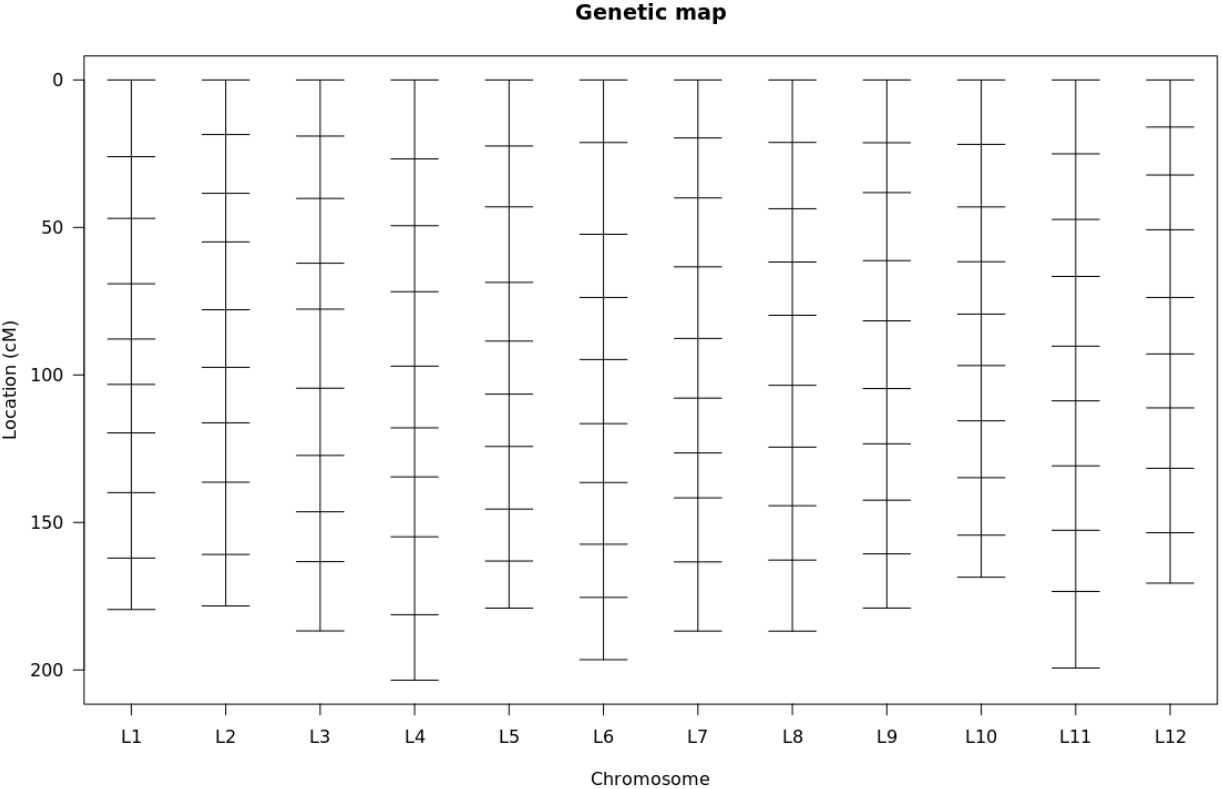
Resulting genetic map produced from ASMap showing the 12 linkage groups and corresponding position of markers in centimorgans

The agreement between simulation parameters and the empirically-determined linkage map values was considered good enough to confirm that the simulator works to model Mendelian segregation correctly. This conclusion supports further use of the simulator to model genetic events in pine populations.

### Simulated effects of inbreeding on growth

Inbreeding significantly affected individual-tree volume in the simulation study (*P* = 4.79e-10) (Table 2). Relative to that of the progeny of outcrosses, volume in the progeny of half-sibs, full-sibs, and selfs was reduced by 4.3, 9.2, and 19.7% (Table 3). Mean volume in the progeny of outcrosses differed significantly from those for all other levels of inbreeding.

**Table 2.** Variance estimates and ratios of estimate and SE for parent and inbreeding by parent interaction for two traits (volume and straightness).

| Effect | Volume |  | Straightness |  |
| --- | --- | --- | --- | --- |
|  | Est | Est/SE | Est | Est/SE |
| Parent | 12.7 | 1.34 | 0.20 | 1.16 |
| Inbreeding x P | 25.35 | 3.31 | 0.13 | 1.39 |

**Table 3.** Mean and percent inbreeding depression at each of the four inbreeding levels for volume and probability of above-average straightness.

| Inbreeding coefficient | Volume (dm<sup>3</sup>) | % ID | Straightness | % ID |
| --- | --- | --- | --- | --- |
| 0 | 124.9 |  | 0.48 |  |
| 0.125 | 119.5 | 4.3 | 0.54 | -12.4 |
| 0.25 | 113.4 | 9.2 | 0.50 | -3.7 |
| 0.5 | 100.3 | 20.0 | 0.50 | -2.9 |

Five parents exhibited a constant decrease in mean volume as inbreeding increased. (Figure 5) Mean volume in two parents showed an increase to the F = 0.125 level but then showed a constant decrease to the F = 0.5 level. In the other three parents, the mean volume of inbred progeny was less than the mean volume in the progeny of outcrosses, with some rank changes among inbreeding levels. The starting number of deleterious alleles within a parent significantly impacted the volume of selfed parents (P=1.14e-4).

**Figure 5.**
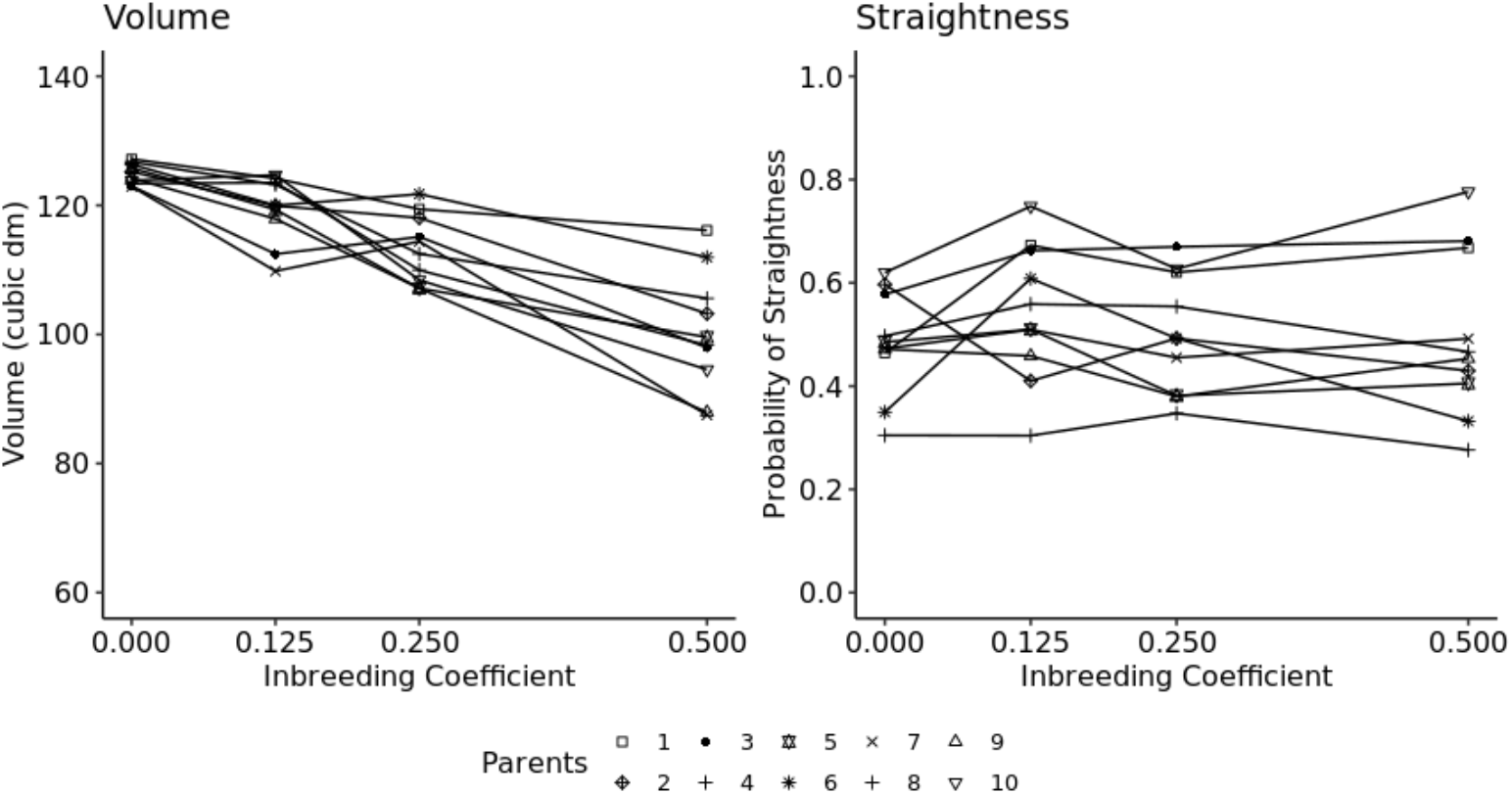
Response of straightness and volume at the four inbreeding levels for ten simulated parents using an inbreeding study design similar to that reported by Ford et al. (2015).

The probability of above-average straightness ranged from 0.48 in the progeny of outcrosses to 0.54 in the progeny of half-sibs (Table 3) and was not significantly affected by inbreeding (*P* = 0.6466) (Table 2). The change in the probability of above-average straightness across the four inbreeding levels was variable in all parents.

The results of the simulated inbreeding study are qualitatively similar to those reported for a study of inbreeding depression in loblolly pine breeding populations (Ford et al, 2015). This suggests that the parameters used to simulate inbreeding depression should provide realistic results for the purposes of comparing breeding population sizes and selection strategies to address the overall goal of the study.

### Effect of selection method on genetic gain and genetic variance

Genetic gain across the ten generations was significantly impacted by the selection method (P=6.3e-3) (Table 4). Across all base population sizes, average genetic values of progeny selections utilizing GBLUP were higher than ABLUP. Regardless of the selection method used, the starting base population size was significant (P=4.78e-8), indicating that a larger base population resulted in more genetic gain. The interaction of method and base population size was also significant (P=2.2e-16), as GBLUP showed a higher rate of genetic gain within a specific base population size than ABLUP. Variance explained by the interaction of selection method and base population with generations were considerable (Est/SE> 2.12).

**Table 4.**
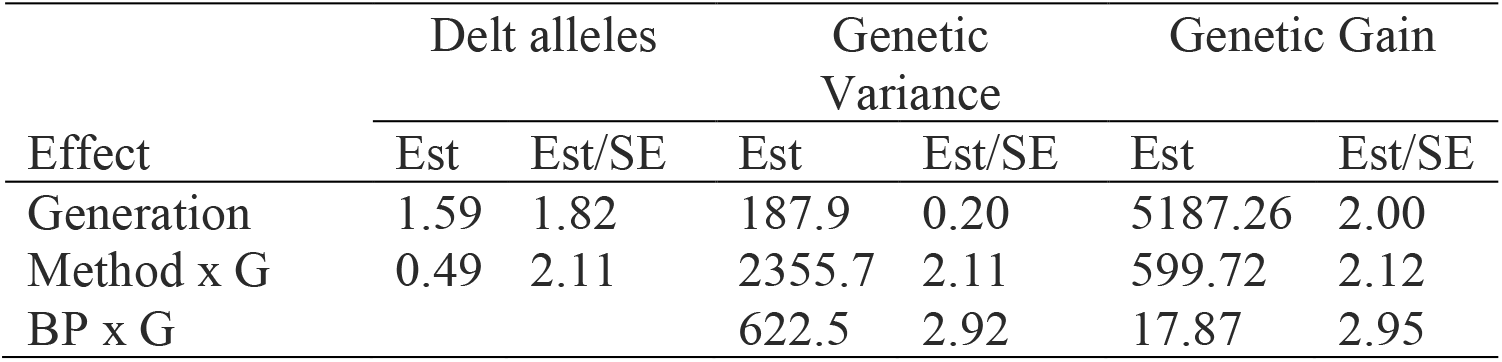
Variance estimates and ratios of estimate and SE for parent and inbreeding by parent interaction for deleterious alleles, genetic variance, and genetic gain.

The impact of the selection method on genetic variance as the main effect was not significant (P=0.054). However, the results of paired t-tests between the genetic variances across base population sizes among the two selection methods ranged from P= 0.969 in generation one to P=2.13e-3 in generation ten, with generations six through ten being significant at the P < 0.05 level. Base population size and the interaction of base population with the selection method used were significant with P=4.23e-3 and P=2.2e-16, respectively. The variance component for generation contributed very little (Est/SE =0.20). In contrast, the interaction of method and generation and base population size and generation contributed significantly (Est/SE> 2.11).

In the first generation, the mean genetic variance of selections was highest in the 280-tree populations and lowest in the 64-tree populations, consistent with the fact that the smaller populations were chosen on the basis of breeding values estimated from simulated open-pollinated progeny tests (Figure 6). The genetic variance for populations selected using ABLUP stayed relatively constant over the last five to six generations of the simulation, while the genetic variance for populations selected using GBLUP declined more sharply starting from the third generation. This difference could result from more efficient within-family selection for genetically-superior offspring, as supported by the dramatically higher rates of genetic gain for all population sizes selected using GBLUP.

**Figure 6.**
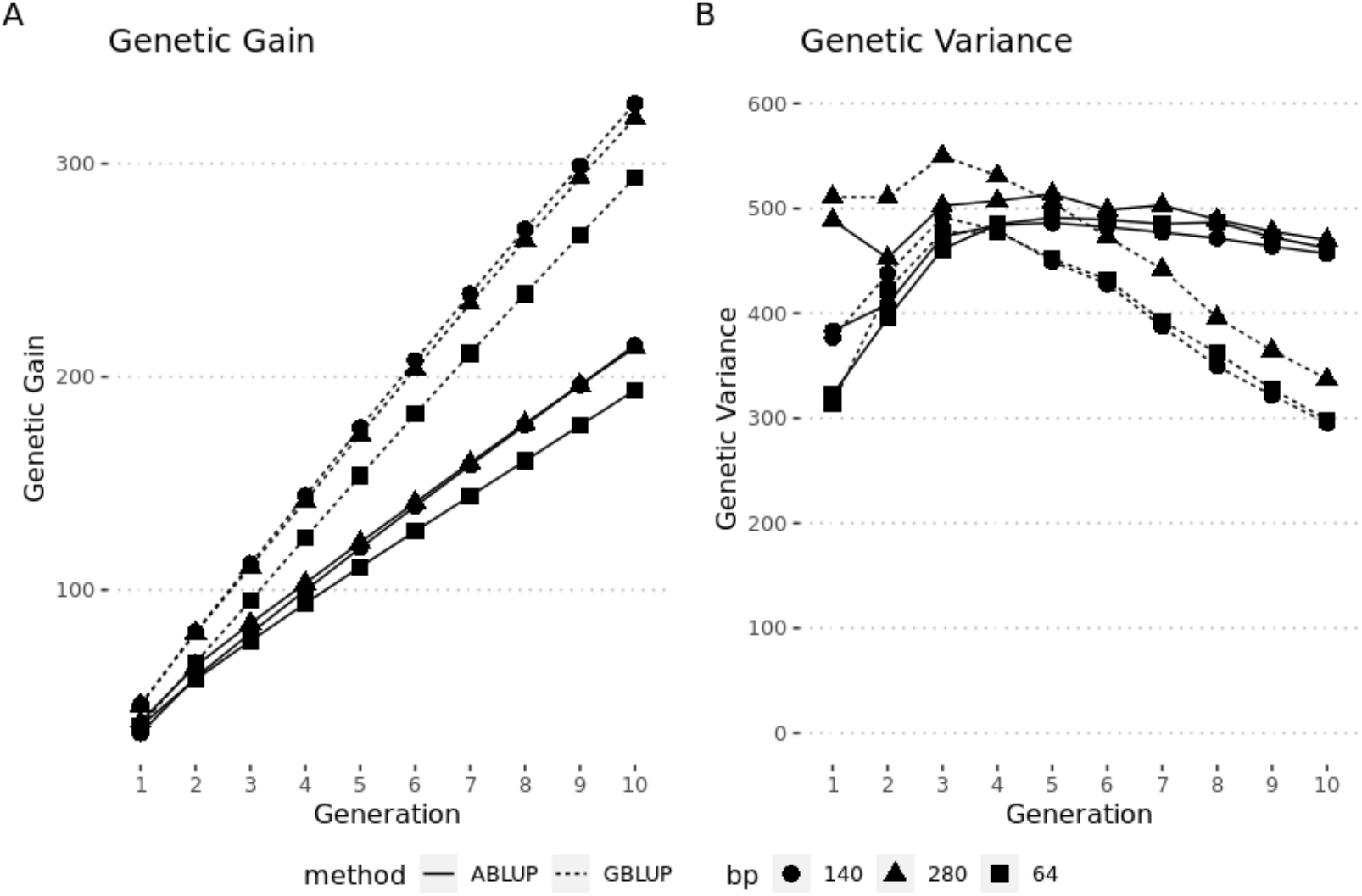
Genetic gain (A) and Genetic Variance (B) of selections made at three different base population levels using two different selection strategies across ten generations

### Effect of selection on the accumulation of deleterious alleles

Method of selection was significant on the number of deleterious alleles over time (P=5.80e-4), with GBLUP removing deleterious alleles at a faster rate than ABLUP. The starting base population size was also significant (P=7.78e-3), with larger starting base populations able to remove deleterious alleles faster. Additionally, the interaction term of base population size and selection method was significant (P=2.49e-9), indicating the rate at which deleterious alleles were purged from the population differed within the base population sizes when using the GBLUP or ABLUP selection method. The Est/SE for the generation variance component was 1.8, contributing relatively small variation to the impact of deleterious allele accumulation. The generation interaction with the method used contributed a moderate amount to variation (Est/SE=2.11), while the interaction of generation and the base population contributed no variation.

Although the average number of deleterious alleles in selections was similar among all strategies in the first generation, selections utilizing GBLUP were consistently lower than ABLUP across the remaining nine generations. (Figure 7). Within GBLUP selection methods, there was no statistical difference between the base populations in the average number of deleterious alleles. In ABLUP, the 64 base population had lower average deleterious alleles than the other base population sizes from generation eight to ten, suggesting that pedigree-based methods purge deleterious alleles more efficiently in smaller breeding populations.

**Figure 7.**
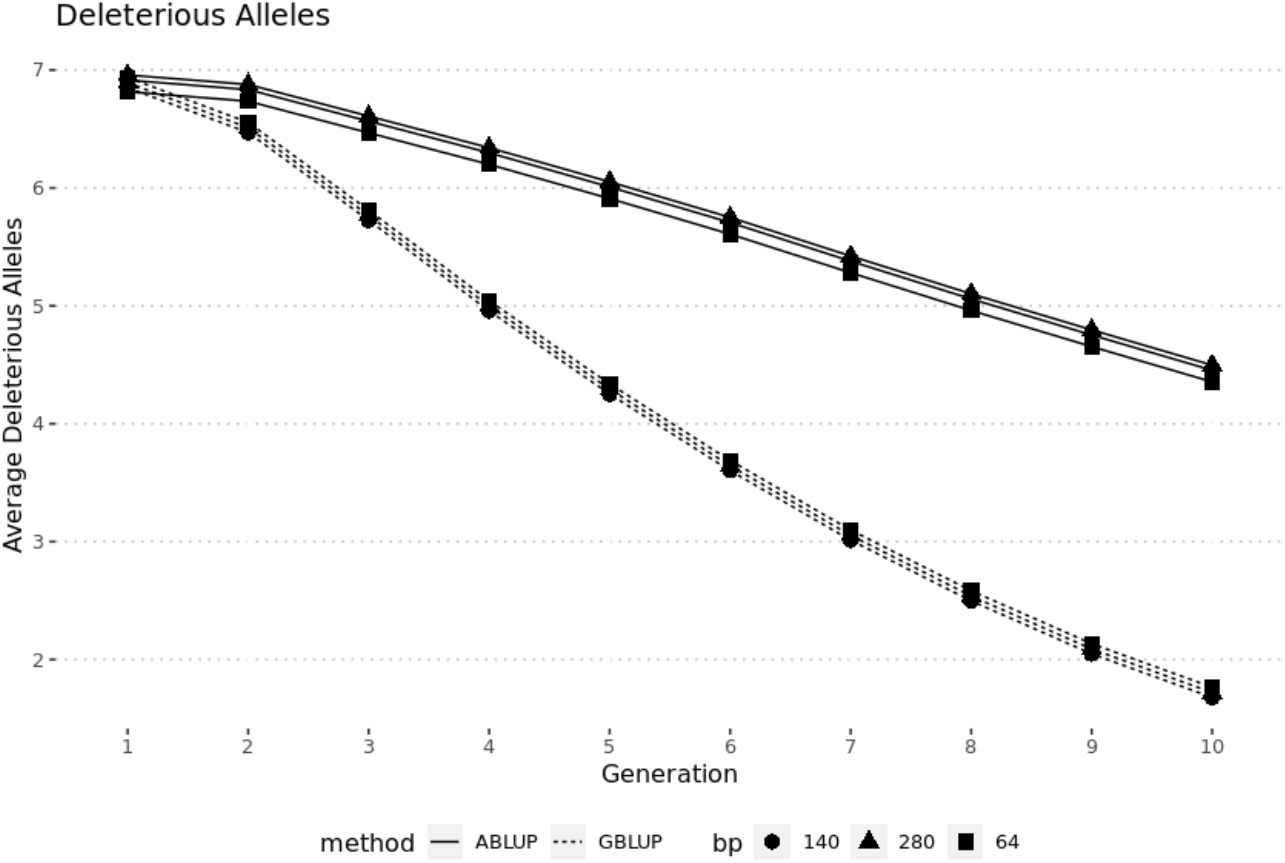
Average number of deleterious alleles within selections made at three different base population levels using two different selection strategies across ten generations

## Discussion

Simulation of breeding strategies for tree species is arguably more critical than for other crops due to the time, cost and space required to conduct multi-generational experiments [31].

Additionally, given the magnitude of inbreeding depression impacts observed in loblolly pine, any breeding method deployed must manage the accumulation of deleterious alleles within the population to ensure short and long-term genetic gain [5-10]. Loblolly pine is a predominantly outcrossing species with a large effective population size [2], and one hypothesis for the cause of observed inbreeding depression is recessive deleterious alleles with significant lethal effects on fitness traits, showing effects only when placed in the homozygous state [47,48]. Contrary to this, smaller populations such as radiata pine are thought to have most of these recessive deleterious alleles purged due to mating among a small remote population [49]. Current efforts are underway to investigate using a bi-allelic SNP marker panel to employ genomic selection in loblolly pine [14].

To explore future opportunities toward managing inbreeding depression while continuing to make genetic gain, we proposed here the idea of utilizing fully informative markers to make selections within-family as opposed to allele-sharing based upon pedigree data. Additionally, we explored maintaining a smaller breeding population with an increase in progeny produced from each cross. A smaller base population, in theory, should be easier to manage, and the increase in the number of progeny per cross should allow for more accurate selections within-family. To accomplish these objectives, we generated a simulation program that can: 1) estimate the total genetic value of individual progeny from simulated crosses, 2) simulate effects of inbreeding depression in progeny after multiple generations of a given breeding strategy, and 3) evaluate ABLUP vs. GBLUP selection strategies utilizing a similar mating design to the one currently employed for their risk of long-term inbreeding depression.

In the simulated inbreeding experiment, similar to the results reported by Ford et al. (2015), inbreeding displayed little impact on the probability of above-average straightness while significantly impacting volume. Although both traits were simulated with many loci having minor additive effects, straightness did not include any loci harboring severe recessive deleterious alleles under dominance control. Modeling the genomic architecture of probability of above-average straightness in this way agrees with the natural breeding history of loblolly pine, in which straightness does not have a severe effect on reproductive success. The results of the simulated inbreeding study agreed with the results of Ford et al. in that all levels of inbreeding depression varied among the parents. This observation could most likely be due to variation among parents in the numbers of lethal and damaging alleles [53,54]. In the simulated inbreeding study, we observed a relationship between the number of recessive deleterious alleles and the effect of inbreeding, and overall patterns of inbreeding agreed with the field study.

Multiple trends were observed among the two selection approaches using pedigree-based (ABLUP) versus marker-based (GBLUP) relationships for within-family selection, regardless of population size. Utilizing GBLUP for within-family selection resulted in a faster reduction of deleterious alleles and a higher rate of genetic gain. The use of markers to make genetic gain has been previously explored and shown to improve selection accuracy compared to pedigree-based relationships, most notably within-family [50,51]. While genomic selection is known to increase coancestry more rapidly and is linked to inbreeding depression, results in this study indicated that even with increased coancestry, we saw deleterious alleles being purged more effectively. We hypothesize that the faster rate of purging deleterious alleles with GBLUP can be attributed to fully informative markers being able to more accurately inform selection decisions that reduce the frequency of deleterious alleles within a family, relative to ABLUP based on pedigree relationships.

The genetic variance from ABLUP for within-family selection was highly variable, whereas when using GBLUP, there was a general decreasing trend over time. Observations that genomic selection increases the loss of genetic diversity as compared to phenotypic selection have been previously shown in an experimental study [55] and stochastic simulations [56,57]. The percentage of genetic variance loss over the ten generations using GBLUP selection was 33.9%, 21.5%, and 4.8% for the 280-tree, 140-tree, and 64-tree base populations. It should be noted, there was still a substantial amount of genetic variance, even after ten generations, which would be approximately 100 years given the current breeding cycle timeline.

Across all base population sizes and selection strategies, there was a trend for purging deleterious alleles and increasing genetic gain over the ten generations. The inference from this observation is that increases in coancestry utilizing either ABLUP or GBLUP will not result in a long-term problem of inbreeding depression. Utilizing an assortative mating design in conjunction with making more crosses than selections seems to be an effective method to potentially expose rare deleterious alleles and remove them from the population.

The current selection method deployed by the NCSU-TIP balances inbreeding depression and genetic gain by maintaining a large base population, and making selections based upon phenotype and average levels of allele sharing extracted from pedigree data [12]. Due to the low individual tree heritability of critical economic traits, accurate estimates of parental breeding values are obtained by producing several progenies for each cross [12]. Within each selection strategy, increasing the base population size from 140 to 280 did not significantly impact genetic gain or genetic variance. The breeding simulations of 280-tree base populations used only half the number of progeny per cross as the 140-tree base population (30 progeny vs 60 progeny) in order to keep the computational requirements of the simulation to a reasonable level. This finding may be attributed to the breeders equation, where gain across generations is a function of the phenotypic heritability of a particular trait, and the force of selection applied [12]. By increasing the number of progeny produced for a given cross and selecting a single individual, phenotypic variance of progeny and the selection intensity applied within family were increased in the 140-tree base population as opposed to the 280-tree base population. Therefore, the comparable rates of genetic gain between the 140-tree population with 60 progeny per cross and the 280-tree population with 30 progeny per cross suggest there may be an advantage to lowering a breeding population size and increasing selection intensity to reduce the workload of managing a large breeding population.

In its current form, the simulator selects individuals for each generation based on breeding/phenotypic values simulated of a single trait. Real breeding programs often work with multiple traits, so the opportunity exists to enhance progeny selection based on multiple traits or a selection index. Future prospective work could also include incorporating other population parameters, such as mutation rate, varying number of markers, numbers of deleterious alleles, dominance coefficients, and allele frequencies, which may help more accurately depict the long-term response of populations to selection.

The overall implications of this study have the potential to have a significant impact on how future selections are made within breeding populations of loblolly pine. All breeding programs aim to increase their ability to make selections with higher accuracy, in less time, with less planting and field experiments. These goals are especially true in loblolly pine due to the slow pace and high cost of breeding and progeny testing. This simulator enables tree improvement programs to model the effects of different user-defined genetic and population parameters on offspring performance and phenotype without the cost or time required for actual breeding experiments. Exploration of breeding strategies through effective simulation could lead to making better selections for future breeding populations, with a higher probability of passing on favorable alleles and providing more overall genetic/economic value to landowners.

## Supporting information

Supplemental Simulator Theory

## References

1. McKeand, S., et al., Deployment of genetically improved loblolly and slash pines in the south. Journal of Forestry, 2003. 101(3): p. 32.

2. Brown, G.R., et al., Nucleotide Diversity and Linkage Disequilibrium in Loblolly Pine. Proceedings of the National Academy of Sciences of the United States of America, 2004. 101(42): p. 15255–15260.

3. Charlesworth, D. and B. Charlesworth, Inbreeding Depression and its Evolutionary Consequences. Annual Review of Ecology and Systematics, 1987. 18(1): p. 237–268.

4. Lynch, M. and B. Walsh, Genetics and analysis of quantitative traits. 1998, Sunderland, Mass: Sinauer.

5. Williams, C.G. and O. Savolainen, Inbreeding Depression in Conifers: Implications for Breeding Strategy. Forest Science, 1996. 42(1): p. 102–117.

6. Halligan, D.L. and P.D. Keightley, How many lethal alleles? Trends in Genetics, 2003. 19(2): p. 57–59.

7. Franklin, E.C., Genetic Load in Loblolly Pine. The American Naturalist, 1972. 106(948): p. 262–265.

8. Remington, D.L. and D.M. O’Malley, Whole-Genome Characterization of Embryonic Stage Inbreeding Depression in a Selfed Loblolly Pine Family. Genetics, 2000. 155(1): p. 337–348.

9. Remington, D.L. and D.M. O’Malley, EVALUATION OF MAJOR GENETIC LOCI CONTRIBUTING TO INBREEDING DEPRESSION FOR SURVIVAL AND EARLY GROWTH IN A SELFED FAMILY OF PINUS TAEDA. Evolution, 2000. 54(5): p. 1580–1589.

10. Ford, G.A., et al., Effects of Inbreeding on Growth and Quality Traits in Loblolly Pine. Forest Science, 2015. 61(3): p. 579–585.

11. McKeand, S.E.; Bridgwater, F.E. 1998. A Strategy for the Third Breeding Cycle of Loblolly Pine in the Southeastern U.S. Silvae Genetica 47, 4(1998)

12. Isik, Fikret & Mckeand, Steven. (2019). Fourth cycle breeding and testing strategy for Pinus taeda in the NC State University Cooperative Tree Improvement Program. Tree Genetics & Genomes. 15. 10.1007/s11295-019-1377-y.

13. Borralho, N.M.G. and G.W. Dutkowski, Comparison of rolling front and discrete generation breeding strategies for trees. Canadian Journal of Forest Research, 1998. 28(7): p. 987–993.

14. Kinghorn, BP, An algorithm for efficient constrained mate selection. Genetics, selection, evolution : GSE, 2011. 43(1): p. 4–4.

15. Isik, F., Genomic selection in forest tree breeding: the concept and an outlook to the future. New Forests, 2014. 45(3): p. 379–401.

16. Queller, DC and KF Goodnight, Estimating Relatedness Using Genetic Markers. Evolution, 1989. 43(2): p. 258–275.

17. VanRaden, P.M., Efficient Methods to Compute Genomic Predictions. Journal of Dairy Science, 2008. 91(11): p. 4414–4423.

18. Meuwissen, T.H.E., B.J. Hayes, and M.E. Goddard, Prediction of Total Genetic Value Using Genome-Wide Dense Marker Maps. Genetics, 2001. 157(4): p. 1819–1829.

19. Bhat Javaid A et al. “Genomic Selection in the Era of Next Generation Sequencing for Complex Traits in Plant Breeding.” Frontiers in genetics vol. 7 221. 27 Dec. 2016, doi:10.3389/fgene.2016.00221

20. Vallejo, R.L., et al., Genomic selection models double the accuracy of predicted breeding values for bacterial cold water disease resistance compared to a traditional pedigree-based model in rainbow trout aquaculture. Genetics, Selection, Evolution, 2017. 49(1): p. 17.

21. Resende, JMFR, et al., Accuracy of genomic selection methods in a standard data set of loblolly pine (Pinus taeda L.). Genetics, 2012. 190(4): p. 1503–1510.

22. Moser, G., et al., A comparison of five methods to predict genomic breeding values of dairy bulls from genome-wide SNP markers. Genetics, selection, evolution : GSE, 2009. 41(1): p. 56–56.

23. Wolc, A., et al., Breeding value prediction for production traits in layer chickens using pedigree or genomic relationships in a reduced animal model. Genetics, selection, evolution : GSE, 2011. 43(1): p. 5–5.

24. Bartholomé, J., et al., Performance of genomic prediction within and across generations in maritime pine. BMC Genomics, 2016. 17(1): p. 604.

25. Tan, B., Grattapaglia, D., Martins, G.S. et al. Evaluating the accuracy of genomic prediction of growth and wood traits in two Eucalyptus species and their F1 hybrids. BMC Plant Biol 17, 110 (2017). https://doi.org/10.1186/s12870-017-1059-6

26. Fu, Y.-B., G. Namkoong, and J.E. Carlson, Comparison of Breeding Strategies for Purging Inbreeding Depression via Simulation. Conservation Biology, 1998. 12(4): p. 856–864.

27. m, v.G., et al., Comparison of two breeding strategies by computer simulation. Crop Science, 2003. 43(5): p. 1764.

28. Rj, K., D. Mj, and T. B, Simulation of the comparative gains from four different hybrid tree breeding strategies. Canadian Journal of Forest Research, 2004. 34(1): p. 209–220.

29. Björklund, M., et al., Assortative mating and the cost of inbreeding — A simulation approach. Ecological Informatics, 2012. 9: p. 59–63.

30. Hallingbäck, H.R., et al., Single versus subdivided population strategies in breeding against an adverse genetic correlation. Tree Genetics & Genomes, 2014. 10(3): p. 605–617.

31. Wu, H.X., et al., Performance of Seven Tree Breeding Strategies Under Conditions of Inbreeding Depression. G3 (Bethesda, Md.), 2016. 6(3): p. 529–540.

32. Godsey, LD, Modeling the financial impact of management decisions on loblolly pine (Pinus taeda) production. 2010, ProQuest Dissertations Publishing.

33. Lin, C.-R.; Buongiorno, J.; Prestemon Jeffrey P.; Skog, K. E. 1998. Growth model for uneven-aged loblolly pine stands : simulations and management implications. (Research paper FPL ; RP-569):13 p. : ill. ; 28 cm.

34. Bishir, J.W. and S. United States. Forest Service. Southern Research, Documentation and user guides for SPBLOB: a computer simulation model of the joint population dynamics for loblolly pine and the southern pine beetle. 2009.

35. R Core Team (2020). R: A language and environment for statistical computing. R Foundation for Statistical Computing, Vienna, Austria. URL https://www.R-project.org

36. Tony Plate and Richard Heiberger (2016). abind: Combine Multidimensional Arrays. R package version 1.4-5. https://CRAN.R-project.org/package=abind

37. Hadley Wickham (2007). Reshaping Data with the reshape Package. Journal of Statistical Software, 21(12), 1–20. URL http://www.jstatsoft.org/v21/i12/.

38. Douglas Bates and Ana Ines Vazquez, (2014). pedigreemm: Pedigree-based mixed-effects models. R package version 0.3-3. https://CRAN.R-project.org/package=.pedigreemm

39. Douglas Bates and Martin Maechler (2015). MatrixModels: Modelling with Sparse And Dense Matrices. R package version 0.4-1. https://CRAN.R-project.org/package=MatrixModels

40. Manos Papadakis, Michail Tsagris, Marios Dimitriadis, Stefanos Fafalios, Ioannis Tsamardinos, Matteo Fasiolo, Giorgos Borboudakis, John Burkardt, Changliang Zou, Kleanthi Lakiotaki and Christina Chatzipantsiou. (2021). Rfast: A Collection of Efficient and Extremely Fast R Functions. R package version 2.0.3. https://CRAN.R-project.org/package=Rfast

41. Julian Taylor, David Butler (2017). R Package ASMap: Efficient Genetic Linkage Map Construction and Diagnosis. Journal of Statistical Software, 79(6), 1–29. doi:10.18637/jss.v079.i06

42. Neves, Leandro Gomide et al. “A high-density gene map of loblolly pine (Pinus taeda L.) based on exome sequence capture genotyping.” G3 (Bethesda, Md.) vol. 4,1 29–37. 10 Jan. 2014, doi:10.1534/g3.113.008714

43. (2008) Haldane’s Mapping Function. In: Encyclopedia of Genetics, Genomics, Proteomics and Informatics. Springer, Dordrecht. https://doi.org/10.1007/978-1-4020-6754-9_7297

44. Wang T, Merkle EC (2018). “merDeriv: Derivative Computations for Linear Mixed Effects Models with Application to Robust Standard Errors.” _Journal of Statistical Software, Code Snippets_,*87*(1), 1–16. doi: 10.18637/jss.v087.c01 (URL: https://doi.org/10.18637/jss.v087.c01).

45. Ott, R. Lyman, and Micheal T. Longnecker. An introduction to statistical methods and data analysis. Cengage Learning, 2015. p. 173–176

46. Douglas Bates, Martin Maechler, Ben Bolker, Steve Walker (2015). Fitting Linear Mixed-Effects Models Using lme4. Journal of Statistical Software, 67(1), 1–48. doi:10.18637/jss.v067.i01.

47. Pollak E. On the theory of partially inbreeding finite populations. I. Partial selfing. Genetics. 1987 Oct;117(2):353–60. doi: 10.1093/genetics/117.2.353. PMID: 3666446; PMCID: PMC1203210.

48. Porcher E, Lande R. Inbreeding depression under mixed outcrossing, self-fertilization and sib-mating. BMC Evol Biol. 2016 May 17;16:105. doi: 10.1186/s12862-016-0668-2. PMID: 27188583; PMCID: PMC4869318.

49. Vogl, Claus, et al. “High resolution analysis of mating systems: inbreeding in natural populations of Pinus radiata.” Journal of Evolutionary Biology 15.3 (2002): 433–439

50. Calleja-Rodriguez, A., Pan, J., Funda, T. et al. Evaluation of the efficiency of genomic versus pedigree predictions for growth and wood quality traits in Scots pine. BMC Genomics 21, 796 (2020). https://doi.org/10.1186/s12864-020-07188-4

51. Velazco, J.G., Malosetti, M., Hunt, C.H. et al. Combining pedigree and genomic information to improve prediction quality: an example in sorghum. Theor Appl Genet 132, 2055–2067 (2019). https://doi.org/10.1007/s00122-019-03337-w

52. Svensson, Julia et al. “Genetic Variation in Height and Volume of Loblolly Pine Open-Pollinated Families During Canopy Closure.” Silvae Genetica 48 (1999): 204–208.

53. Bramlett D.L. Pepper W.D.. 1974. Seed yield from a diallel cross in Virginia pine. P. 49–55 in Proc. of a colloquium: Seed yield from southern pine seed orchards. Georgia Forest Research Council, Macon, GA.

54. Skrøppa T. 1996. Diallel crosses in Picea abies. II. Performance and inbreeding depression of selfed line. For. Genet. 3(2): 69–79.

55. Rutkoski, J., et al. “Genetic gain from phenotypic and genomic selection for quantitative resistance to stem rust of wheat.” (2015).

56. Jannink, Jean-Luc. “Dynamics of long-term genomic selection.” Genetics Selection Evolution 42.1 (2010): 1–11.

57. Lin, Z., et al. “Genetic gain and inbreeding from genomic selection in a simulated commercial breeding program for perennial ryegrass. Plant.” Genome 9 (2016).

