## Supplemental Simulator Theory for "Simulation of Pedigree vs. fully-informative marker based relationships matrices in a Loblolly Pine breeding population"

**SIMULATION OF A GENETIC MAP (MAKE_GENETIC_MAP)** 2

**DETERMINING SIZE OF LINKAGE GROUPS** 2
**TYPES OF LOCI** 3
**DISTRIBUTION OF LOCI** 3
**INTERVALS AND RECOMBINATION RATES** 3
**MINOR ALLELE FREQUENCIES** 4

**CREATION OF FOUNDER POPULATIONS (CREATE_PARENTS)** 5

**ALLELIC VALUES** 5
*INBRED PARENTS 5
**ASSIGNMENT OF ALLELES** 6
*HETEROZYGOUS MARKERS 6
*MODELING UNRELATED FOUNDERS OF AN OUTCROSSING SPECIES 6

**GENERATION OF CROSS DESIGN (CREATE_CROSS_DESIGN)** 7

**GENERAL MATING DESIGNS** 7
CROSS FILE INPUT 7
SELF 8
RANDOM MATING 8
HALF-DIALLEL 8
**RANKED MATING DESIGNS** 8
CO-ANCESTRY THRESHOLD 8
RANKING SCHEME 8
SINGLE-PAIR 9
ASSORTATIVE MATING 9
**OUTPUT** 10

**SIMULATED RECOMBINATION (MAKE_CROSSES)** 12

**CALCULATION OF GENETIC VALUES (CALC_TGV)** 14

**SIMULATION OF PHENOTYPES (SIM_PHENOS)** 16

***PROGENY SELECTION* (EXTRACT_SELECTIONS)** 17

**SELECTION STRATEGY** 18
PHENOTYPES 18
GBLUP 18
ABLUP 20
**OUTPUT** 20
INBREEDING ESTIMATES 21
PRODUCING RELATIONSHIP MATRIX OF SELECTIONS 22

**OPEN POLLINATION TESTING (OP_TESTING)** 23

### Simulation of a genetic map (make_genetic_map)

***User required inputs***

| | |
|---|---|
| *num.lgs* | Number of linkage groups |
| *map.length* | Total genetic map length in centimorgans |
| *num.markers* | Number of genetic makers |
| *total.qtl* | Total number of quantitative trait loci |
| *num.snpqtl* | Number of quantitative SNP loci |
| *map.dist* | Character string indicating one of either "haldane" or "kosambi" to specify which method will be used to calculate genetic map distances |

***Optional inputs*** *(see details)*

| | |
|---|---|
| *distribute.loci* | Changes distribution of loci on linkage groups |
| *marker.distribution* | Changes distribution of markers place on genetic map |
| *linkage.size.range* | Changes the linkage group size range |
| *snp.qtl.maf* | A vector of two numbers indicating the min and max frequency of SNP QTL minor alleles |
| *loci.per.lg* | Specify number of loci to assign to each linkage group |
| *marker.maf* | Vector to specify the minor allele frequency for each marker |

```
create_genetic_map( num.lgs=12, map.length=1800 , num.markers=120,  num.snpqtl=1960,
total.qtl = 2640, snp.qtl.maf = c(.01,.02))
```

#### Determining size of linkage groups

The first step in creating a genetic map is determining the length of each individual linkage group by sampling random deviates from a uniform distribution with a mean ($\mu$) equal to the total map length in centimorgans ($ML_{cM}$) divided by the total number of linkage groups ($LGs$). The maximum and minimum of the uniform distribution is the $\mu$ +/- the linkage size range ($lsr$) multiplied by the $\mu$, respectively.

$$\mu = \frac{ML_{cM}}{\#\ of\ LGs}\ , min = \mu - lsr * \mu\ , max = \mu + lsr * \mu$$

### Types of Loci

Once the linkage group sizes are determined, three types of loci: markers, SNP QTL, and random QTL (rQTL) may be distributed among the linkage groups as specified by the input parameters: *num.markers*, *total.qtl*, and *num.snpqtl*. The *total.qtl* parameter is interpreted as including the number of both random and SNP QTL. Random QTL represent loci which may contribute random genetic noise, possibly by events such as epistasis or regulatory interactions, while SNP QTL represent loci which may be set to display dominance effects. Any amount remaining from *total.qtl* – *snp.qtl* will act as rQTL so that the total number of loci to be distributed among the linkage groups is equal to the number of markers plus the total number of QTL.

### Distribution of Loci

By default, the number of loci placed on each linkage group are distributed randomly. Loci may be distributed among the linkage groups evenly so that each has the same number of loci using *distribute.loci = "even"*. The distribution may be provided by the user as a list format with *distribute.loci = "list"*.

The markers and QTL (both SNP and random) are distributed randomly across the map. In place of random assignment, markers may also be distributed evenly along each linkage group by setting *marker.distribution = "equally-spaced"*. If markers are set to be equally-spaced, the process of selecting which loci will be markers is determined using the following format:

For each linkage group, identify the true average distance ($AD_{LG}$) that a single marker ($M_i$) would be from the previous marker by dividing the last loci position for that linkage group ($LP_{LG}$) by the total number of markers ($M_{LG}$).

$$AD_{LG} = \frac{LP_{LG}}{M_{LG}}$$

Multiply each $M_i$ by $AD_{LG}$ to get a vector of what the "perfect locus position" would be if markers were equally spaced. Then, for each marker identify a locus position on the genetic map which minimizes the absolute difference between the "perfect locus position" and all available loci positions.

### Intervals and recombination rates

In all the above cases, number of intervals to be estimated for the genetic map is equal to the total number of loci minus 1. The intervals, in Morgans, between each locus and the next are determined by random sampling from a uniform distribution with:

$$min = 0.2 * \frac{linkage.size}{\#loci}, max = 1.8 * \frac{linkage.size}{\#loci}$$

For each linkage group, simulated intervals between loci are used to calculate recombination rates by using either the Haldane or Kosambi mapping function, as specified by the *map.dist* parameter [1, 2]. After interval distances are used to estimate recombination frequencies, they are converted back to cM and output in the resulting genetic map as loci positions along each linkage group.

### Minor Allele Frequencies

By default, markers and random QTL are sampled from random deviates of a beta distribution with $\alpha = .4$ and $\beta = .4$. Marker MAFs may be changed to a user provided distribution or fixed number through the *marker.distribution* parameter. The SNP QTL MAFs are sampled from a uniform distribution with a min and max that is set by the user using the *snp.qtl.maf* input.

The resulting output of the function is a matrix with 7 columns:

| LG | locus | dist | types | maf | pos | recfreqs |
|---|---|---|---|---|---|---|
| Linkage group number | Locus name | Interval distance | rQTL, snp, or marker | Minor Allele Frequency | Position along the Linkage Group | Recombination frequencies |

### Creation of founder populations (create_parents)

***User required inputs***

| | |
|---|---|
| *map.info* | Genetic map matrix returned from make_genetic_map( ) |
| *num.parents* | A number indicating how many parents are to be created |

***Optional inputs***

| | |
|---|---|
| *max.delt.alleles* | The maximum number of deleterious alleles any single parent can have. Default: 0 |
| *par.markers.unique* | Logical. Should each parent have unique alleles as markers? Default: FALSE |
| *heterozygous.markers* | Logical. Should all marker loci be heterozygous Default: FALSE |
| *inbred.parents* | Logical. If TRUE with par.markers.unique set to TRUE will produce inbreds. Default: FALSE |
| *QTL.sd* | A number providing the standard deviation of random QTL effects generated for the parents. Default: 0.25 |

```
create_parents(map.info=the.map, num.parents=96, max.delt.allele=14, par.markers.unique=T)
```

#### Allelic values

Alleles assigned are either characters, in the case of SNP QTL and marker loci, or numbers in the case of random rQTL. By default, the markers and SNP QTL are assigned the characters "a" and "c" to represent the major and minor allele, respectively. The value assigned to markers may be changed to be unique in every individual by setting *par.markers.unique* equal to TRUE, in which case, multiple characters will represent the major and minor allele of each individual so that no two parents have similar markers (e.g. Parent 1: "AA" and "aa"; Parent 2: "AB" and "ab"). The rQTL are always assigned numbers sampled from the addition of two normal distributions with a mean of 0 and standard deviation half of that specified by the *QTL.sd* variable. The default *QTL.sd* is set to 0.25 making the possible allelic values of rQTL to be 1,0, or -1.

##### *Inbred Parents

Setting *inbred.parents* to equal TRUE will follow the same general rules as above with the exception of markers. Since the purpose of this simulator was not directly intended for inbred crops this setting will create inbred parents who each have unique identical homozygous markers at each allele e.g. (Inbred 1 = "A" "A", Inbred 2 = "b" "b")

### Assignment of alleles

**By default, assignment of major and minor alleles for marker and SNP QTL is done by testing the genetic map MAFs against a set of randomly sampled numbers from a uniform distribution ranging from 0 to 1. Each parent is assigned two alleles at every locus. For each of the two alleles, an independent sample of random numbers is generated and tested against the map MAFs. Loci which have a MAF less than that of the random number sampled are assigned to have a minor allele and loci greater than the number sampled are assigned to have the major allele. Given that the rQTL are numeric, one value to represent each allele at a single locus is sampled from the distribution previously mentioned.

***Special Cases:*

#### *Heterozygous markers

Setting *heterozygous.markers* equal to TRUE samples half of the loci and assigns allele 1 to be the minor allele. The remaining half of loci for allele 1 that was not sampled is assigned to have the major allele. The second allele is assigned so that all loci which were previously assigned the minor allele are assigned the major and vice versa.

#### *Modeling unrelated founders of an outcrossing species

For our purposes, the *mean.delt.alelle* parameter was included so that we may model long-term effects of highly deleterious loci. By indicating that the maximum number of deleterious alleles is > 0 the SNP QTL are treated as loci which harbor highly deleterious alleles. The impact this has is that SNP QTL in the founder population will be generated so that no single individual shares a minor allele at a given locus. Additionally, no individual will have two minor alleles at any one locus. This occurs through a loop format with vectors that keep track of which positions are available. It is important to note that if there are not enough SNP QTL specified to assign in this fashion, the function will not work.

The number set through *mean.delt.allele* is interpreted as the mean number of deleterious alleles that may be in any single individual within the population. It is also used to sample the number of deleterious alleles that will be assigned to each founder parent. For each of the two alleles of every individual, the number of deleterious alleles is determined by sampling from a range of 0 to the *mean.delt.allele*. No individual may have a total number of deleterious alleles greater than 2**mean.delt.allele* in the founder population.

The resulting output of the function is a list object that contains the following:

| Output Name | Info |
|---|---|
| ***delt.allele*** | The number of recessive SNP QTL in each founder parent |
| ***genos.3d*** | 3-dimensional array containing genotypes of parents |
| ***num.parents*** | Value indicating the total number of parents generated |
| ***parent.IDs*** | Vector containing the name of parents |
| ***parent.Marker.matrix*** | A matrix containing Parent markers |
| ***parent.SNPQTL.matrix*** | A matrix containing the Parent SNP QTL character (“a” or “c”) |

### Generation of cross design (create_cross_design)

***User required inputs***

| | |
|---|---|
| *parent.info* | Object returned from either the create_parents() or extract_selections() |
| *prog.percross* | Number of progeny that should be made per cross |
| *generation* | Number that specifies which generation |
| *mating.design* | Specify the type of mating design to be used in making crosses: "AM" (Default), "Self", "RM", "SP", "HD", or "cross.file.input". (See details for description) |

***Optional inputs***

| | |
|---|---|
| *coancest.thresh* | Logical. Should a co-ancestry threshold be used as to minimize making related crosses? |
| *use.op.par.phenos* | Logical. Set to TRUE if passing parents created from op_testing() |
| *cross.file* | User provided cross file. Should be used if mating.design = "cross.design.input" |
| *parent.phenos* | Object returned from sim_phenos() |
| *use.par.marker.effects* | Logical. Set to TRUE if passing parents created from op_testing() |

```
create_cross_design(parent.info=the.parents, prog.percross=60, generation=1,
mating.design="RandomMating")
```

In general, mating designs may be broken into two categories: general and ranked mating designs. The difference between the two is general mating designs do not consider relatedness among parents or progeny, whereas ranked mating designs can utilize a co-ancestry threshold and prioritize crosses based on estimated breeding values or phenotypes. The four general mating designs include selfing, half-diallel, random mating, or the input of a user-specified cross file. Ranked mating designs include single-pair and assortative mating. In all mating designs, the generation number and data object returned from one of *create_parents(), extract_selections(),* or *op_testing()* is required.

#### General Mating Designs

##### Cross file input

To input a cross design, the source location of a cross file as .txt or .csv may be input to the *cross.file* parameter with *mating.design* set to equal “cross.file.input”. The file should contain three columns (no column names necessary) and rows which indicate Parent 1, Parent2, and number of progeny. The names of parents within the text file should correspond to the column names of the *genos.3d* object that is included in the *parent.info* parameter.

#### Self

Setting *mating.design* equal to "Self" will generate a cross design that replicates all the column names in the *genos.3d* object as Parent 1 and Parent 2. The number specified in *prog.percross* will be used to determine how many individuals will be generated for each self.

#### Random Mating

Specifying random mating ("RM") will sample half of the column names from the *genos.3d* object and place them as Parent 1, while the unselected half will be placed as Parent 2. The number specified in *prog.percross* will be used to determine how many individuals will be generated.

#### Half-Diallel

A half-diallel ("HD") mating design generates a cross file in which all unique pairwise crosses are made between individuals (column names of the *genos.3d* object). The number specified in *prog.percross* will be used to determine how many individuals will be generated.

### Ranked Mating designs

#### Co-ancestry threshold

A co-ancestry threshold is intended to be used with ranked mating designs when trying to minimize breeding among similar individuals while attempting to maximize genetic gain. Since the founder population is assumed to be completely unrelated, using the *coancest.thresh* parameter only has an effect starting at generation 2 after running *extract_selections()*. If *coancest.thresh* is set to TRUE, a relationship matrix among selections from the previous generation, created from either pedigree or marker data, is used to screen which crosses can be made with respect to a certain co-ancestry threshold.

The threshold begins at 0.01 and continues up to 1 by 0.0005. At each cutoff, pairwise column and row names from the relationship matrix which have a value less than the threshold are subset using the reshape2 package and the ranked mating design specified is attempted to be created **[CITATION].** If the requirements for a mating design cannot be met at the first threshold, the number is raised by 0.005 iteratively until requirements may be met. The final level of co-ancestry that was applied to the cross design is output by the function and stored as a list variable named *coancestry.threshold*.

#### Ranking scheme

In all ranked mating designs, genetic gain is attempted to be maximized by emphasizing the crossing of individuals which have the highest estimated breeding values (EBVs) or phenotypes. Since we assume no selection has occurred before the first generation, utilizing one of these mating designs with the founder population will result in parents being ranked by phenotypes created using *sim_phenos()*. In the case of progeny, EBVs are estimated during *extract_selections()* by using the user-specified method of selection (ABLUP, GBLUP, or Phenotype; see *extract_selections*). If one of these mating designs is used with co-ancestry threshold set to TRUE, the ranking method above is the same except for that the only crosses which pass the co-ancestry threshold will be ranked and potentially used.

#### Single-Pair

Specifying a single-pair ("SP") mating design outputs a cross file whose total number of crosses is equal to ½ the number of parents or previous selections. Therefore, if using this method there must be an even number of individuals in the previous generation from which the mating design is constructed. In the case of no applied co-ancestry threshold, the phenotypes or EBVs of prospective parents are first sorted from highest to lowest. Then from the sorted vector, every other individual is selected from 1 to *n individuals* to be Parent 1 and the remaining are designated as Parent 2. Assigning parents in this fashion ensures that each parent will only be present in a single-pair. An emphasis on genetic gain is made by mating the two individuals with the highest breeding values.

When a co-ancestry threshold is applied to this mating design, the ranking and assignment of parents works the same as above, with the exception that all crosses made must have a co-ancestry value as determined from the relationship matrix that is less than that of the current threshold. If this requirement is not met due to some of the suggested crosses being too similar, the level of the co-ancestry threshold will increase until it the requirements of the mating design are met.

#### Assortative Mating

An assortative mating ("AM") design is implemented by first sub-setting the total number of individuals into four unique groups (Q1, Q2, Q3, and Q4) depending upon their EBVs or phenotypes. The four groups represent four quantiles so that if the number of parents was 16, Q1 would include individuals 1-4 (the top 25% of sorted individuals), Q2 would be individuals 5-8 (top 26-50%), Q3 would contain individuals 9-12 (lower 51-75%), and Q4 would hold the remaining individuals 13-16 (bottom 76-100%).

If using an assortative mating scheme without a co-ancestry threshold, the groups are used to assign crosses in the following way:

- Each of the parents in Q1 (top 25%) are used a minimum of 4 times:
    - Two crosses are generated for each parent in Q1 by assigning $parent_i$ as the first parent and randomly selecting two other individuals within Q1 to be the second parent.
    - One cross is generated between a randomly selected $parent_i$ in Q1 and randomly selected individual from Q2.
    - One cross is generated between a randomly selected $parent_i$ in Q1 and a single randomly selected individual from Q3
- Individuals in Q2 are used in three total crosses
    - The first time as mentioned above with Q1
    - A second time by randomly sampling parents without replacement from Q2 and Q3
    - Finally, a third time by sampling a random parent without replacement from Q2 and Q4

The resulting cross design file ultimately has a minimum of four crosses with individuals from Q1, three crosses with individuals from Q2, two crosses with individuals from Q3, and one cross containing individuals from Q4. Designing the mating design in this way places an emphasis on breeding the top individuals while attempting to maintain diversity by including a low number of crosses with the lower ranked individuals.

If this method is used with a co-ancestry threshold, the assignment of individuals which will be crossed is the same as above except for crosses made among the Q1 group (Q1xQ1) and between the Q1xQ2 groups. For each of those two types of crosses, relatedness estimates subset from the relationship matrix must be less than the co-ancestry threshold. Starting with the top individual from Q1 possible crosses which satisfy the threshold are sorted by mid-parent mean and the top prospective cross is utilized. It is important to note that while each parent in Q1 is still used a minimum of 2 times when crossing to another individual in Q1 and once when crossing to an individual from Q2, unlike the non-co-ancestry method, the other parent selected is based on mid-parent mean calculation instead of being randomly sampled.

For Q1xQ1 this means that although reciprocal crosses are not included, if an individual in Q1 is not related to any of the other individuals, it will be used as parent 2 more often since the resulting mid-parent mean would be the highest. With respect to crosses made between Q1xQ2 all parents must have at least one cross that is less than the provided threshold. Starting with the top parent from Q1, the relationship matrix is subset to include any prospective crosses which pass the co-ancestry threshold. The resulting crosses are then sorted by mid-parent mean and the top potential cross is made. The second top parent from Q1 must then also have a prospective cross which passes the co-ancestry threshold and does not include parent 2 that was selected in the first cross. This pattern continues until all parents from Q1 and Q2 have a single cross.

### Output

In addition to generating a cross design file, this function is responsible for keeping track of the pedigree and progeny/parent names. The assignment of names to new progeny is done on a continuous scale so that if, for example, individuals to be mated have the names "1" to "64" the first few progenies from the first row of the cross design will be named "65", "66", "67", etc. The naming of progeny this way ensures that a pedigree will not have duplicate names of individuals from past generations. All outputs are listed below:

| Output Name | Info |
|---|---|
| ***coancestry.threshold*** | Value indicating the co-ancestry threshold if coancest.threshold was set to TRUE |
| ***cross.design*** | Cross design matrix that will be used to generate progeny |
| ***full.pedigree*** | Pedigree of all individuals |
| ***num.crosses*** | Value indicating the number of crosses which will be made |
| ***num.parents*** | Value indicating the number of unique parents in the cross design |
| ***parent.IDs*** | Vector containing the names of progeny which will be created |
| ***progeny.pedigree*** | Pedigree matrix containing only parents used to create the cross design and progeny to be created |
| ***selection.ped*** | Pedigree matrix including only past selections |
| ***total.progeny.number*** | Value indicating the total number of progeny that will be generated |

### Simulated recombination (make_crosses)

***User required inputs***

| | |
|---|---|
| *parent.info* | Object returned from create_parents() or extract_selections() |
| *map.info* | Object returned from create_genetic_map() |
| *cross.design* | Object returned from create_cross_design() |

***Optional inputs***

| | |
|---|---|
| *run.parallel* | Logical. Set TRUE to run function in parallel |
| *num.cores* | The number of cores to use if running in parallel |

```
make_crosses(parent.info=the.parents, map.info=the.map, cross.design=cross.file,
run.parallel=T, num.cores=3)
```

Implementing this function requires the R package abind and can be multi-threaded through using the parallel package by setting *run.parallel* equal to TRUE and specifying a digit for the number of cores to utilize in the *num.cores* input **[CITATIONS].** Whether the function is ran using multi-threading or not, the methods for generating progeny and simulating recombination in all crosses are the same. Genotypes of individuals within the cross file who will be mated are stored in the object returned from either the *create_parents()* or *extract_selections()* function.

The creation of progeny among the individuals in the cross file is done by generating two independent gametes, one from each parent, that become allele 1 and allele 2 for a given progeny. Recombination within each respective gamete is simulated so that each progeny receives a single unique gamete from each parent of a cross.

The following is an example of how one unique gamete is assigned from a single parent to progeny:

- First, columns in the genotype array which correspond to the names of parent 1 and parent 2 for a cross are subset so that allele 1 and 2 from each respective parent are in separate matrices. The first gamete assigned to a progeny is from parent 1 and the second gamete is from parent 2.

- For each linkage group, gamete 1 is simulated by initially sampling a set of numbers from 0 to 1 by .0001 that is equal to the length of loci on that respective linkage group. Then, a logical test is performed between the sampled numbers and the recombination frequencies on the genetic map for loci which correspond to a given linkage group. If the recombination frequency on the genetic map is greater than the sampled number, a recombination will occur at that locus. If no recombination frequencies on the map are greater than the sampled number for a given locus, a new sample is taken so that at least one recombination occurs for every linkage group. The locations of recombination for each linkage group are then used to assign the final gamete from a parent to a progeny.

- For each linkage group, the number 1 or 2 is sampled to indicate whether the assignment of alleles to the progeny will begin with parent allele 1 or parent allele 2. If 1 is sampled, progeny will be assigned the alleles of Parent 1 allele 1 until a recombination takes place. At loci where recombination occurs, the allele assigned to progeny switches from Parent 1 allele 1 to Parent 1 allele 2. The same process occurs for the second gamete that is from Parent 2. Each progeny is created independently so that different recombination patterns occur within progeny from a single cross.

The output of this function is a 3-dimensional array that contains progeny genotype values and has dimensions equal to *n loci x n progeny x 2*.

### Calculation of genetic values (calc_TGV)

***User required inputs***

| | |
|---|---|
| *map.info* | Genetic map matrix returned from create_genetic_map( ) |
| *geno.info* | Object containing genotype information. Should be from create_parents() or extract_selections() |
| *A* | Value assigned to the Major SNP QTL allele |
| *a* | Value assigned to the minor SNP QTL allele |

***Optional inputs***

| | |
|---|---|
| *founder* | Logical. Set to TRUE if determining genetic values for a founder population. (Default: FALSE) |
| *dom.coeff* | Value indicating the dominance coefficient applied to SNP QTL alleles |
| *par.markers.unique* | Logical. Set to TRUE if parents have unique markers (Default: FALSE) |
| *rqtl.dom* | A vector specifying dominance values for any rqtl loci |

```
calc_TGV(geno.info=the.parents, map.info=the.map, A=1, a=-100, dom.coeff=1,founder=T)
```

The primary purpose of this function is to calculate the genetic values of individual parents or progeny produced with either *create_parents()* or *make_crosses()*, respectively. Note that if using on individual parents it is important to set *founder* equal to true so that the parent names can be obtained. All genetic values are calculated from an individuals' random QTL (rQTL) and SNP QTL.

The rQTL, which range from -1 to 1, are summed across all loci with respect to a given individual.

$$rQTL.effects_i = \sum_{1}^{n} rQTL_n$$

Loci which are SNP QTL are handled according to the value assigned to the major allele (*A*), minor allele (*a*), and level of dominance (*dom.coeff*). If the dominance coefficient assigned is 0, the SNP QTL will be additive so that the contribution of SNP QTL to an $Individual_i$ genetic value is:

$$SNP\ QTL\ effects = (num.major.alleles_i * A) + (num.minor.alleles_i * a)$$

If non-zero is assigned to dominance, the contribution of SNP QTL is calculated so that loci which are homozygous for the major allele will be equal to $A$, loci homozygous for the minor allele will be equal to $a$, and those which are heterozygous loci will be $A$*dom.coeff.

$$SNP\ QTL\ effects = (num.AA_i * A) + (num.aa_i * a) + \ (num.Aa_i * A) + \ (num.aA_i * a)$$

The final genetic value of $Individual_i$ is then determined by the sum of all its rQTL and SNP QTL effects:

$$Individual_i = rQTL.effects_i + SNP.QTL.effects_i$$

In addition to a vector of genetic values, this function will return a bi-allelic SNP marker matrix and corresponding marker map. The marker matrix has dimensions equal to *n loci* * *n progeny* and is coded as 0, 1, or 2 indicating the number of minor alleles an individual has at a given locus. The marker map returned includes the linkage group name and position of each marker along the genetic map.

| Output Name | Info |
|---|---|
| ***genetic.values*** | Vector containing the genetic values of progeny |
| ***marker.loci*** | Vector indicating the rows of the genetic map which are markers |
| ***marker.map*** | Data frame containing linkage group and position information of markers |
| ***markers.matrix*** | Matrix containing bi-allelic SNP markers of progeny |
| ***snp.effect.values*** | Vector containing the summed value of SNP QTL effects for each individual |

### Simulation of phenotypes (sim_phenos)

***User required inputs***

| | |
|---|---|
| *TGV.object* | Object returned from calc_TGV() function |
| *h2* | Value specifying the individual tree-heritability |

***Optional inputs***

| | |
|---|---|
| *E.var* | Value indicating a specific environmental variance |

sim_phenos(TGV.object=parent.TGV, h2=.3)

Phenotypes of individuals may be simulated utilizing genetic values returned from *calc_TGV()*. The total variance of all genetic values ($\sigma_G^2$) and the heritability specified are utilized to simulate environmental noise. The standard deviation used in sampling random deviates is used based on the following assumptions:

The individual tree-heritability can be calculated as: $h^2 = \frac{\sigma_G^2}{\sigma_P^2}$

Therefore: $\sigma_G^2 = h^2 * \sigma_P^2$

Solving for the phenotypic variance: $\sigma_P^2 = \frac{\sigma_G^2}{h^2}$

Since total phenotypic variance is determined by: $\sigma_P^2 = \sigma_G^2 + \sigma_E^2$

The environmental variance can be calculated as: $\sigma_E^2 = \sigma_P^2 - \sigma_G^2$

Solving for $\sigma_E^2$: $\sigma_E^2 = \frac{\sigma_G^2}{h^2} - \sigma_G^2$

Therefore, random deviates from a normal distribution were sampled using the following mean and standard deviation:

$$\mu = 0 \ , \ \sigma = \sqrt{\frac{\sigma_G^2}{h^2} - \sigma_G^2}$$

A single deviate is sampled for each $Individual_i$ and added to the genetic value to be output as phenotypes:

$$Individual_i = sampled.deviate_i + genetic.value_i$$

Instead of sampling environmental noise in this way, a standard deviation of which to sample from can be input as a value through the *E.var* parameter. The output of this function includes simulated phenotypes and corresponding genetic values for each individual.

### *Progeny Selection* (extract_selections)

***User required inputs***

| | |
|---|---|
| *map.info* | Genetic map matrix returned from create_genetic_map( ) |
| *cross.design* | Object returned from create_cross_design() |
| *past.tgv* | Object returned from calc_TGV of parents in cross design |
| *past.phenos* | Object returned from sim_phenos of parents in cross design |
| *parent.info* | Object returned from either create_parents() or previous extract_selections() |
| *progeny.info* | Object returned from make_crosses() |
| *progeny.TGV* | Object returned from calc_TGV of progeny in cross design |
| *progeny.phenos* | Object returned from sim_phenos of progeny in cross design |
| *relationship.matrix.type* | Either "pedigree" or "markers" indicating the type of relationship matrix to use for selection and next generation cross design |
| *numselections.within.family* | Value indicating how many selections should be made within each family |
| *num.selections.among.family* | Value indicating how many selections should be made among families |
| *among.family.selection* | One of "Phenotype", "ABLUP" or "GBLUP" indicating method to make selections on breeding values |
| *within.family.selection* | One of "Phenotype", "ABLUP" or "GBLUP" indicating method to make selections on breeding values |

***Optional inputs***

| | |
|---|---|
| *run.parallel* | Logical. Set TRUE to run function in parallel |
| *num.cores* | Value indicating the number of cores to use |
| *weighted* | Logical. Should genomic-relationship matrix be weighted by pedigree-relationship matrix? Default: FALSE |

| | |
|---|---|
| *the.weight* | Value indicating the weight to multiply by pedigree-based relationship matrix if weighted is TRUE. |
| *reduced* | Logical. Should be set to TRUE if calling function using subset of parents returned from op_testing(). Default: FALSE |

```
extract_selections( map.info=the.map, cross.design=cross.object,  progeny.info=progeny1, parent.info=the.parents, ,progeny.TGV=progeny1.TGV, progeny.phenos=progeny1.PHENOS, num.selections.among.family=64, num.selections.within.family=1, relationship.matrix.type="markers", selection.strategy="GBLUP", past.tgv=parent.TGV, past.phenos=parent.PHENOS)
```

The selection of individuals is accomplished using the *extract_selections()* function and requires the R packages: MatrixModels, parallel, pedigreemm, and pedigree. There are three main methods for evaluating, ranking, and making forward selections among progeny: 1) Phenotype, 2) pedigree-based best linear unbiased predictions (ABLUPs), or 3) genomic-based BLUPs (GBLUPs). Within each of the three methods, the number of selections that are made among and within families for a given generation are user-specified. After selections are made, a relationship matrix composed of either pedigree or markers is generated from selections and can be used with the co-ancestry method in *create_cross_design()* for the next generation. Additional stats such as the inbreeding coefficient of the selection pedigree, number of deleterious alleles, or genetic and phenotypic gain are also output by the function.

### Selection strategy

#### Phenotypes

Selecting phenotypes as the selection strategy will use progeny phenotypes created from the *sim_phenos()* function to rank and select individuals. For each family, the mean phenotype of progeny is calculated and families are sorted in decreasing order. The number of among family selections (*n*) is then used to subset the top *n families*. Within each of the top *n families*, the number of specified selections for within family (*z*) is used to select *z progeny* from that family.

#### GBLUP

The GBLUP selection strategy utilizes a progeny genomic-relationship matrix and their corresponding simulated phenotypes to estimate breeding value predictions (EBVs). The genomic-relationship matrix may be weighted by the pedigree-based matrix if *weighted* is set to TRUE and a value is present in the *weight* parameter. For example, if *weighted* equals TRUE and *the.weight* is set to 0.01 then the relationship matrix used to estimate BLUPs will be equal to:

$$Relationship\ matrix = Genomic_{RelMat} * 0.99 + Pedigree_{RelMat} * 0.01$$

When using the pedigree-relationship to weight the genomic-relationship matrix, the selection pedigree is created using *getA()*from the pedigreemm package. Progeny which correspond to those in the current generation are then subset from the pedigree relationship matrix so that it can be multiplied by the weight factor and added to the genomic relationship matrix.

#### *Creation of genomic relationship matrix*

If markers are bi-allelic so that the major allele is "a" and minor allele is "c", the genomic-relationship matrix is created using *calcG()* from the pedigree package. However, if markers are created in parents to be multi-allelic so that they are unique to every individual in the founder population, a relationship matrix is constructed by shared character string matching among each of the progeny.

An example of estimating relatedness between two progenies using their first and second allele for a set of five markers:

| ***Marker locus #*** | **P1 A1** | **P1 A2** | **P2 B1** | **P2 B2** |
|---|---|---|---|---|
| *M1* | AA | bb | bb | AA |
| *M2* | aa | BB | BB | AA |
| *M3* | aa | bb | aa | BB |
| *M4* | AA | AA | AA | AA |
| *M5* | AA | AA | BB | BB |

For each marker, a single individual (*P1*) with two alleles is matched to the each of the other progeny alleles (*P2*) in six different possible ways:

- *A1.1* = Allele A1 matches B1 and B2
  - M4
- *A1.2* = Allele A1 only matches B1
  - M3
- *A1.3* = Allele A1 only matches B2
  - M1
- *A2.1* = Allele A2 matches B1 and B2
  - M4
- *A2.2* = Allele A2 only matches B1
  - M1, M2
- *A2.3* = Allele A2 only matches B2
  - None

Using each of the six vectors a 2,1, or 0 is assigned to each marker depending on if the two progenies share both alleles, one allele, or no alleles.

- Share two alleles:
  - Loci which are in both A1.1 and A2.1: M4
  - Loci which are in A1.2 & A2.3: None
  - Loci which are in A1.3 & A2.2: M1
- Share one matching allele:
  - Loci only in A1.1, A1.2, A1.3, A2.1, A2.2, or A2.3: M2, M3
- Share no matching alleles:

- Loci which are not in any of the above categories: M5

Since two perfectly identical individuals would share *x alleles* equal to 2* *n markers*, the number (2,1, or 0) assigned to each marker may be added and divided by the total possible shared alleles to get percentage similarity:

- Assigned 2: M1, M4
- Assigned 1: M2, M3
- Assigned 0: M5
- (2*2 + 1*2 + 0) / 10 = 60% similarity between progeny 1 and progeny 2

The percentage similarity between the two individuals is then placed in the cell of the relationship matrix which corresponds to the row of progeny 1, column of progeny2. The same method is utilized for all progeny until the relationship matrix is complete.

#### *Estimation of BLUPs*

Once a genomic-relationship matrix is calculated, and weighted by the pedigree-relationship matrix if applicable, the inverse of the matrix is calculated and progeny BLUPs are estimated in the following way:

```
n.col <- ncol(the.data)
h.2 <- var(prog.genetic.values)/var(progeny.phenos)
lambda <- (1-h.2)/h.2
I <- diag(n.col)
sol <- solve(rBind(cBind(n.col, t(rep(1,n.col))),
            cBind(rep(1,n.col), (I + (lambda*the.data)))),
        matrix(c(sum(progeny.phenos),
              c(as.vector(progeny.phenos)))))
```

The BLUPs returned for progeny are then used for selection in the same format as Phenotypes using the number of specified among and within family selections.

#### ABLUP

Pedigree based BLUP estimates of progeny are calculated by first obtaining the A inverse of the selection pedigree (all previous selections and progeny of current generation) using *getAInv()* from the pedigree package. The rows and columns of the A inverse matrix which correspond to progeny of the current generation are then subset from the matrix and used in the same fashion as in the GBLUP approach. Additionally, selections that will be made from each family is identical to the approach used in either the Phenotypes or GBLUP selection strategy.

### Output

The names of selections which are made using one of the above strategies are then subset from multiple objects that contain information about the progeny (e.g. genotypes, phenotypes, pedigree) and merged with previous parent information. In addition, information extracted from selections is used to generate various stats on things such as the level of inbreeding among selections or the change in genetic variance due to selection. The following table may be referenced to see all outputs of this function:

| Output Name | Info |
|---|---|
| ***all.genetic.vals*** | Vector containing genetic values of all previous selections (including founder parents) |
| ***all.markers*** | Matrix containing bi-allelic SNP markers of progeny and past parents |
| ***all.phenos*** | Vector containing all phenotypes of past and current selections |
| ***bulmer.effect*** | Value representing the change in total progeny genetic variance due to selection |
| ***delt.alleles*** | Vector containing the number of minor allele SNP QTL in each selected individual |
| ***fullped*** | A pedigree matrix containing the complete pedigree |
| ***genos.3d*** | 3-Dimensional array containing genotype information of current and previous selections |
| ***num.parents*** | Value indicating the number of selections made in the current generation that may be used as parents for the next generation |
| ***prog.inbred.level*** | Vector containing the inbreeding coefficient for all progeny |
| ***relmat*** | A relationship matrix of selections that will be used if co-ancestery threshold is set to TRUE in create_cross_design() of the next generation |
| ***select.EBVs*** | Vector containing the estimated breeding values of selections returned from selection strategy |
| ***select.genval*** | Vector of genetic values corresponding to selections made |
| ***select.inbred.level*** | Vector containing the inbreeding coefficient for only selected progeny |
| ***select.ped.ids*** | Vector containing names of selections which were made |
| ***selection.ped*** | A pedigree matrix including all previous selections |
| ***selection.phenos*** | Vector containing the phenotypes of selected individuals |

The genetic values of selections are returned in the resulting object as the name *select.genvals* and may be used to estimate the mean or variance of the selected progeny values. The output *bulmer.effect* is calculated as the change in the variance of genetic values due to selection:

$$bulmer.effect = \sigma^2_{G(selections)} - \sigma^2_{G(all_{progeny})}$$

Additionally, phenotypes of the selected individuals are returned in the *selection.phenos* output and EBVs of progeny estimated using one of the selection strategies are returned as *select.ebvs*. If using one of the ranked cross design methods in the next generation, the selection EBVs will be used to rank prospective parents and crosses.

### Inbreeding estimates

The inbreeding coefficient of the entire pedigree (all individuals from all generations whether they were selected or not) is estimated using the inbreeding function from the pedigree package. The inbreeding coefficients of all progeny from the current generation are then subset from the resulting vector and represent the *prog.inbred.level*. Similarly, the progeny which were selected from the current generation are subset from the resulting vector and represent the *select.inbred.level*. In the case of modeling deleterious alleles, information on the number of minor SNP QTL alleles that are present within each of the selected individuals may be obtained in the *delt.alleles* output. The numbers represented in the output for each selection are generated by identifying how many "c" alleles each selection as at its' respective SNP QTL loci.

### Producing relationship matrix of selections

A relationship matrix of selections may be constructed using either the pedigree or markers depending on how *relationship.matrix.type* was set when calling the function. The construction of either relationship matrix works the same as it does in the GBLUP selection strategy. A pedigree based relationship matrix is constructed from the selection pedigree utilizing *getA()*and a marker-based relationship matrix is created using either *calcG()* for bi-allelic or shared character matching for multi-allelic. The relationship matrix created will be used in the next generation when creating a cross design if using the co-ancestry threshold set to TRUE.

### Open Pollination Testing (op_testing)

***User required inputs***

| | |
|---|---|
| *map.info* | Genetic map matrix returned from make_genetic_map( ) |
| *parent.info* | Object returned from create_parents |
| *parents.phenos* | Object returned from sim_phenos |
| *parents.TGV* | Object returned from calc_TGV |
| *num.select* | Value specifying the top number of parents to subset |
| *cross.prog* | Value indicating the number of progeny to make for each cross |
| *A* | Value assigned to the Major SNP QTL allele |
| *a* | Value assigned to the minor SNP QTL allele |
| *h2* | Value specifying the individual tree-heritability |

***Optional inputs***

| | |
|---|---|
| *num.cores* | Value indicating the number of cores to use |
| *run.parallel* | Logical. Set TRUE to run function in parallel |
| *E.var* | Value indicating a specific environmental variance |
| *dom.coeff* | Value indicating the dominance coefficient applied to SNP QTL alleles |

This is a wrapper function that simulates open pollination among a set of founders. The function requires that a genetic map and parents are previously generated using *make_genetic_map()* and *create_parents()*. The first step includes the creation of a cross design where all individuals are mated to every other individual, replicating a full-diallel mating design. The number of progeny that are simulated for each cross is determined by the *cross.prog* parameter. For each cross simulated recombination among parents, calculation of progeny genetic values, and estimation of progeny phenotypes is identical to that in *make_crosses()*, *calc_TGV()*, and *sim_phenos()* respectively. The mean of phenotypes that are simulated for progeny from each OP parent is taken so that parents may be ranked from best to worst with respect to its' general combining ability with other individuals in the population. Once founder parents are ordered, the *num.select* parameter can be used to subset the top *n parents* from the founder population. The output of this function contains multiple objects (see below) and may be used as an input to *create_cross_design()* to generate a specified mating design only using a subset of parents ranked by OP family mean phenotypes.

| Output Name | Info |
|---|---|
| ***delt.alleles*** | Vector containing the number of minor allele SNP QTL in each selected individual |
| ***genetic.values*** | Vector containing genetic values of parents |
| ***genos.3d*** | 3-dimensional array containing genotypes of parents |
| ***marker.matrix*** | Matrix containing bi-allelic SNP markers (see calc_TGV) |
| ***pars*** | Vector containing the sorted OP family mean phenotypes |
| ***phenos*** | Vector containing phenotypes of parents |
